# Idiosyncratic choice bias in decision tasks naturally emerges from intrinsic stochasticity in neuronal network dynamics

**DOI:** 10.1101/284877

**Authors:** Lior Lebovich, Ran Darshan, Yoni Lavi, David Hansel, Yonatan Loewenstein

## Abstract

Idiosyncratic tendency to choose one alternative over others in the absence of an identified reason is a common observation in two-alternative forced-choice experiments. It is tempting to account for it as resulting from the (unknown) participant-specific history and thus treat it as a measurement noise. Here we quantify idiosyncratic choice biases in a perceptual discrimination task and a motor task. We report substantial and significant biases in both cases that cannot be accounted for by the experimental context. Then, we present theoretical evidence that even in idealized experiments, in which the settings are symmetric, idiosyncratic choice bias is expected to emerge from the dynamics of competing neuronal networks. We thus argue that idiosyncratic choice bias reflects the microscopic dynamics of choice and therefore is virtually inevitable in any comparison or decision task.

---

Decision making is the cognitive process of choosing an action among a set of alternatives. Decision making is often studied in experiments, composed of trials, each associated with a single decision. While a decision in a trial is primarily determined by the relevant features of the alternatives in that trial, biases are commonly observed^1^. Of specific relevance to this work are participant-specific tendencies to prefer one alternative over the other(s). Such biases, which we term idiosyncratic choice biases (ICBs) have been described as early as half a century ago in perceptual discrimination^2–4^ and operant learning tasks^5–7^.

In discrimination tasks, the ICBs interfere with the estimate of perceptual noise. In operant learning experiments these biases mask the learning behavior. That is why such biases are typically considered as nuisance. When analyzing choice behavior, these biases are often accounted for by adding an ad-hoc participant-specific bias parameter^3^ or by counterbalancing the choices to average them out.

Many factors can contribute to ICBs. For example, in perceptual discrimination tasks, a stimulus in a given trial is often perceived as being more similar to the stimuli presented in previous trials^8–10^. Similarly, participants tend to choose those actions that were previously more often rewarded^11,12^. Finally, participants may exhibit a preference towards an alternative because the corresponding motor action requires the least effort. Heterogeneity between the participants along any of these factors is sufficient to generate ICBs. One may thus expect that these biases would be diminished if these factors are controlled for in the experimental design or are factored out in the analysis. In contrast to this expectation, here we argue that even in an idealized gedanken experiment, in which symmetry between subjects in all the above factors is kept, substantial ICBs are expected. These ICBs that cannot be accounted for by the experimental context are the subject of this study.

We quantify ICBs in a perceptual discrimination task and in a novel sensory-motor task, in which sequential and operant factors are controlled for. We then analyze the ICBs in the framework of a Drift Diffusion Model (DDM) and show that they are primarily the result of biased drift rates. Finally, we show analytically and numerically that ICBs naturally emerge from the intrinsic stochasticity of the dynamics of competing populations of spiking neurons. Our work thus suggests that ICBs are inevitable unless they are actively suppressed, e.g. by the reward schedule.

## Results

### ICBs in the bisection discrimination task

We quantified ICBs in the bisection discrimination task depicted in Fig. 1a (inset). In each trial, a vertical transected line was presented on the screen and participants were instructed to indicate the offset direction of the transecting line (see Materials and Methods). Fig. 1a depicts the fraction of an ‘Up’ response, *p*_up_ as a function of the offset for three participants. As expected, the probability of a correct response increased with the magnitude of the offset *ΔL* ≡ (*L*^*U*^ − *L*^*D*^)/(*L*^*U*^ + *L*^*D*)^, where *LU* and *L*^*D*^ denote the lengths of the Up and Down segments of the vertical line. However, the responses differed between the three participants: the blue psychometric curve is shifted to the right of the black curve, whereas the red curve is shifted to its left.

**Figure 1:**
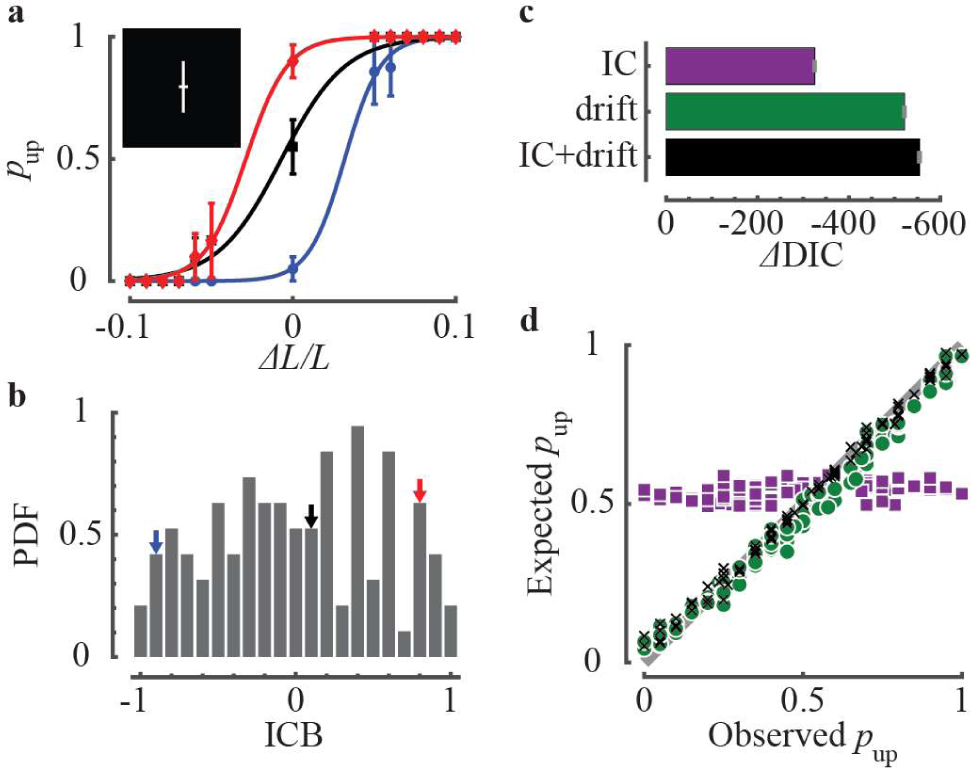
ICBs in the vertical bisection task. **a**, Psychometric curves of three participants: the observed fraction of responding ‘Up’, *p*_up_, vs. the sensory offset *ΔL*/*L*. Error bars denote the standard error of the mean (SEM). Curves are best-fit logistic functions. Inset, a schematic illustration of the stimulus in a single trial. **b**, Distribution of ICBs (ICB = *p*_up_ − *p*_down_) of all participants (*n* = 100). The ICBs of the three participants in (b) are denoted in the histogram by arrows of corresponding colors. **c**, Model comparison using DIC (Materials and Methods). The DIC of the ‘IC bias’ DDM (purple), ‘drift bias’ DDM (green) and ‘IC+drift bias’ DDM (black) were measured relative to the baseline DDM, ΔDIC = DIC_model_ − DIC_baseline_. Error bars are SEM, based on three repetitions of the fitting procedures. Results indicate that the ‘IC+drift’ DDM accounts for the data slightly better than the ‘drift bias’ DDM and much better than the ‘IC bias’ DDM. d, Relative contributions of the drift-bias and IC-bias to the ICBs in the ‘IC+drift bias’ DDM. Each symbol depicts a single participant. Abscissa: the observed *p*_up_ in the impossible trials. Ordinate: expected *p*_up_, based on average posteriors of each participant in the biased ‘IC+drift’ DDM (Eq. 6 in Materials and Methods). Black Xs: both initial conditions and drifts were taken from the ‘IC+drift’ DDM. Green circles: Drifts were taken from the ‘IC+drift’ DDM with symmetric initial conditions (*z* = 0.5 in Eq. 6). Purple squares: Initial conditions were taken from the ‘IC+drift’ DDM assuming no drift (*A* = 0 in Eq. 6). Gray line is the diagonal. Slopes of best-fit orthogonal regressions are: black Xs, 0.96; green circles, 0.93; purple squares, 0.04.

We considered the choices of the participants in 20 “impossible” trials (1/6 of the trials), in which the line was transected at its midpoint (*ΔL* = 0). The participant whose psychometric curve is plotted in black in Fig. 1a responded ‘Up’ in 11/20 impossible trials, which is statistically indistinguishable from chance (p=0.82, two-sided Binomial test). By contrast, the two other participants (red and blue in Fig. 1a) exhibited significant choice biases, responding ‘Up’ in 18/20 and 1/20 of the trials, respectively (p<0.001, two-sided Binomial test). Overall, 48% of the participants (n=100) exhibited a significant choice bias (24% significant ‘Up’, p < 0.05, two-sided Binomial test; 24% significant ‘Down’, p < 0.05, two-sided Binomial test). These ICBs were not restricted to the impossible trials. Rather, they were also observed in the possible trials albeit to a lesser degree. Biases in the possible and impossible trials were highly correlated (Fig. S1; two-sided Pearson’s *ρ*=0.63, p<10^-12^). At the population level, we could not detect a global bias. The fraction of ‘Up’ choices in the impossible trials across all participants was 0.505, which is not significantly different from chance (95% CI 0.45-0.56, bootstrap).

To quantify the heterogeneity of these ICBs across the population, we computed for each participant the difference between the fraction of ‘Up’ and ‘Down’ responses in the impossible trials. This measure quantifies the bias because it vanishes for unbiased choices (ICB = 0 for *p*_up_ = 0.5) and its magnitude is maximal if choices are deterministic (ICB = −1 for *p*_up_ = 0; ICB = 1 for *p*_up_ = 1). The distribution of ICBs across the participants is depicted in Fig. 1b. Its width is a measure of idiosyncrasy of these biases across the participants. We found that the variance of the distribution is significantly larger than expected by chance (p<10^-6^, one-sided bootstrap test, Bernoulli process). These results further establish the existence of ICBs in the vertical bisection task.

As mentioned in the Introduction, operant effects can contribute to ICBs. To minimize the contribution of feedback to the ICBs, participants received only sparse feedback every 30 trials on their accumulated performance until that point. Another potential contributor to ICBs is a propensity to repeat in a trial the actions taken in the previous trials. To minimize sequential effects, the impossible trials were always preceded by three irrelevant trials (see Materials and Methods). Indeed, the probability that a participant would repeat in an impossible trial the action she took in the previous (possible) trial was 0.50 ± 0.01 (average over participants ± SEM) (see also Fig. S2).

We then analyzed the ICBs in the framework of the DDM. According to the DDM, noisy evidence in favor of each alternative is integrated over the course of the trial. The difference of these evidence, a quantity known as the decision variable, is computed and a decision is reached once this variable reaches one of two decision thresholds. The DDM has been extensively used to explain both behavioral and neurophysiological data^13–18^. In this framework, ICBs in the impossible trials can emerge via two mechanisms. In the first mechanism, the bias results from the initial condition of the decision variable being not equidistant from the two thresholds. In the second mechanism, the bias results from a drift bias of the decision variable, which is unrelated to the veridical evidence^19–22^.

We investigated which of these two mechanisms best accounts for the ICBs which we observed experimentally. To that end, we fit choices and reaction-times of participants in the impossible trials to four versions of the DDM. The goodness of each fit was assessed using the Deviance Information Criterion (DIC; Materials and Methods). The first model was a baseline DDM with symmetric, i.e., equidistant initial condition and no drift bias. By construction, there are no ICBs in this model and it was only used as a baseline for comparison with the other three models. To dissect the relative contributions of the drift and initial condition to the ICBs, we added to this baseline DDM (1) idiosyncratic drift rates (‘drift bias’ DDM), (2) idiosyncratic initial conditions (‘IC bias’ DDM) or (3) both idiosyncratic drift rates and idiosyncratic initial conditions (‘IC+drift bias’ DDM). The DICs of all three models were compared to the DIC of the baseline model. As depicted in Fig. 1c, all three models did better than the baseline model. The ‘drift bias’ DDM (green) did substantially better than the ‘IC bias’ DDM (purple). These results suggest that in the framework of the DDM, bias in initial condition contributes less to the observed ICBs than the bias in the drift rate. We further dissected the relative contributions of the drift and IC to the ICBs in the ‘IC+drift’ DDM (black), which did better than the other models (Fig. 1c). To that goal, we computed for each participant the ICB expected from the DDM with parameters extracted from the ‘IC+drift’ DDM. As shown in Fig. 1d (black Xs), the expected and observed ICBs are in good agreement. They are also in good agreement when instead of the extracted initial conditions, symmetric ones are used (green circles). This indicates that asymmetry in the initial conditions does not play an important role in the generation of the ICBs. Indeed, when using the extracted initial conditions but unbiased drift we failed to account for the ICBs (purple squares).

### ICB in the motor task

Next, we constructed a novel motor task, in which ICBs are unlikely to emerge from idiosyncratic sensory asymmetries. In each trial, two adjacent colored dots were displayed on a white circular background (Inset in Fig. 2a, see also Fig. S3a). Participants were instructed to drag, as fast as possible, these two dots into a central region indicated by a larger black disk. To ensure that the participants would make two temporally-separated reaching movements, we introduced a 1.1 sec delay after the completion of the dragging of the first colored dot (Materials and Methods). The task was presented to the participants as a motor-speed task, in which faster movements are more rewarded (see Materials and Methods). However, the behavioral parameter that we were interested in was the order in which participants chose to execute the two dragging movements. In that sense, this paradigm is a symmetric, binary, implicit, decision-making task. In this task, choice bias manifests as a particular preference in the order in which the two dots are dragged. Each participant was presented with 10 pairs of dots, each pair differing in colors and locations. Each of these pairs was presented 20 times in a pseudorandom order. The ICB of a participant for a given pair of dots was defined as the difference between the fraction of trials in which the clockwise (CW) and counterclockwise (CCW) dot was dragged first (ICB = *p*_cw_ − *p*_ccw_). This allowed us to measure 10 different ICBs (one for each pair) for each participant.

**Figure 2:**
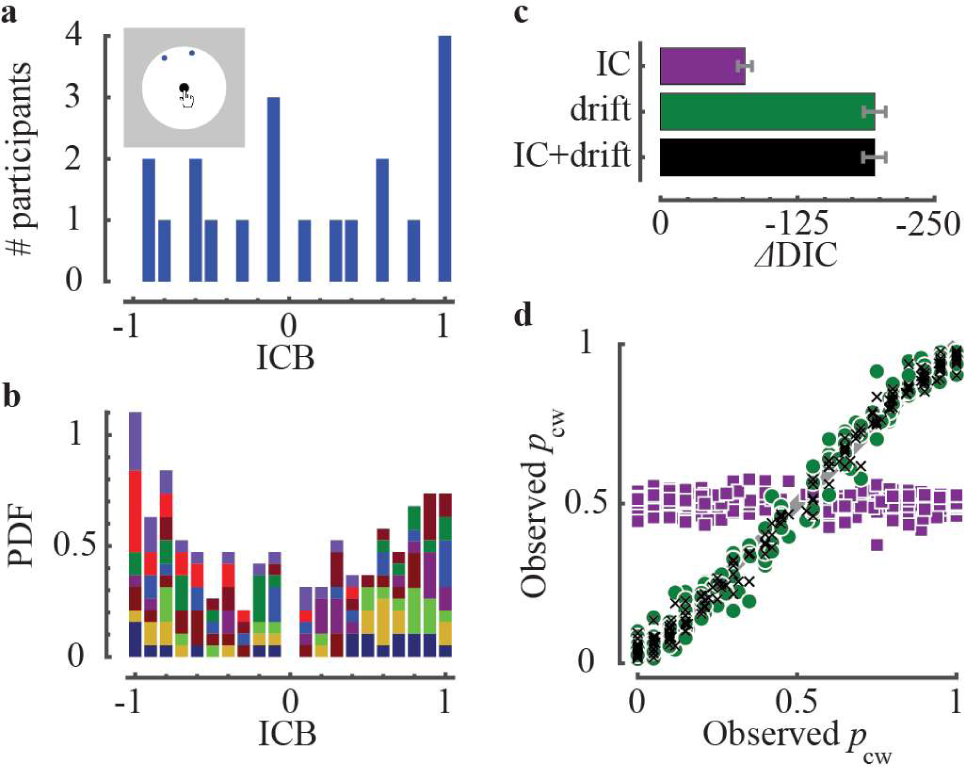
ICBs in the motor task. ICB = *p*_cw_ − *p*_ccw_ where *p*_cw_ and *p*_ccw_, are the probabilities of choosing first the clockwise and the counter clockwise dot. **a**, The distribution of ICBs of all participants (*n* = 20) for the pair of dots in the inset. **b**, The distribution of ICBs for all 10 pairs of dots in the experiment (color-coded as in Fig. S3b). **c**, Model comparison using DIC, as Fig. 1c. Model fits were performed separately on each pair of dots. Bars and error bars denote the average and SEM, *Δ*DIC, over the 10 pairs of dots. **d**, Same as Fig. 1d, demonstrating that the ICBs in the motor task are dominated by the drift-biases. Slopes of best-fit orthogonal regressions are: black Xs, 0.98; green circles, 0.99; purple squares, −0.01.

Figure 2a depicts the distribution of choice biases across the participants for a particular pair of dots (inset). At the population level, we could not detect a global bias. The fraction of clock-wise choices across all participants was 0.55, which is not significantly different from chance (95% CI 0.40-0.70, bootstrap). Nevertheless, 65% of the participants exhibited significant ICB for this pair (35% significant preference towards choosing first the clock-wise dot and 30% significant preference in favor of choosing first the counter-clockwise dot; p < 0.05 two-sided Binomial test). Consistent with that, the variance of the distribution of preferences in that pair was significantly larger than expected by the population-average preference (p<10^-6^, one-sided bootstrap test, Bernoulli process). Variance of the distributions of preferences that is significantly larger than expected by the population-average preference was observed in all ten pairs (Fig. S3b).

We then analyzed the ICBs using the DDM framework. As in the bisection task, the ‘drift bias’ model (green) did substantially better than the ‘IC bias’ model (purple) for all ten pairs, indicating a smaller contribution to behavior of the biased initial conditions relative to the contribution of biased drift rates. The DIC of the ‘IC+drift bias’ DDM (black) was comparable to the DIC of the ‘drift bias’ DDM (Fig. 2c). In half of the pairs, the DIC of the ‘IC+drift bias’ DDM model was the lowest, whereas in the other half, the DIC of the ‘drift bias’ DDM was the lowest. We further dissected the contributions of drift biases and initial conditions to the observed ICBs using the ‘IC+drift bias’ DDM (Fig. 2d). As in the bisection task, we found that drift bias, rather than asymmetry in the initial conditions, is the dominant contributor to ICBs in the motor task.

### ICBs in the Poisson network model

What underlies participant-to-participant differences in drift rates in the bisection and motor tasks? To address this question, we constructed a simple neuronal network model of decision making and used it to study behavior in the bisection task. It consists of two populations of neurons representing ‘Up’ and ‘Down’ choices, denoted by ‘U’ and ‘D’ (Fig. 3a, left). Each population is made of *N*/2 independent Poisson neurons, such that the spike train of each neuron in a trial is an independent homogeneous Poisson process. The firing rates of the neurons depend on the offset in the input (*ΔL*) such that the firing rates of the U neurons increase with *ΔL*, whereas that of the D neurons decrease with *ΔL* (Fig. 3a, right). In addition, each neuron receives an offset-independent input, which captures the heterogeneity in the firing rates of the neurons within each population (see Eq. 1 in Materials and Methods). Specifically, the firing rates of the neurons are drawn from log-normal distributions, whose parameters depend on the offset (orange and pink distributions in Fig. 3a). In the absence of an offset (*ΔL* = 0), the firing rate distributions of the two populations are the same (blue distribution in Fig. 3a, right). Both the Poisson-like firing of action potentials^23^ and the log-normal distribution of firing rates^24,25^ are hallmarks of cortical dynamics.

**Figure 3:**
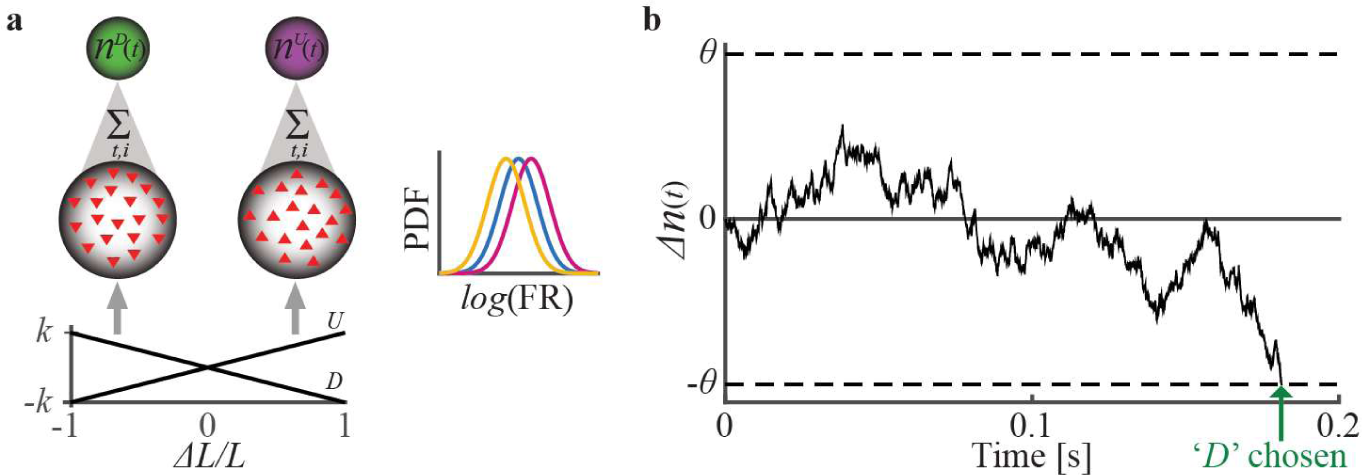
The Poisson network model. **a**, Schematic illustration of the network. It consists of 2 populations of independent Poisson neurons (Center), receiving stimulus-selective input (Bottom). Direction of triangle denotes selectivity to the visual offset, *ΔL*, as in Fig. 1. The neurons emit spikes, which are accumulated (Top). Right, the stimulus-dependent distribution of firing rates. In the absence of offset (*ΔL* = 0), the rate of ‘*U*’ and ‘*D*’ neurons are drawn from the same distribution (blue curve). When the upper segment of the line is the longer (*ΔL* > 0), neurons in population ‘*U*’ increase their firing rates (pink curve), whereas neurons in population ‘D’ decrease their firing rates (orange curve). Note that the lognormal distribution of rates is equivalent to normal distribution of log-rates. **b**, Example trial. The absolute value of the difference in spike counts is accumulated over time, until the threshold is reached. The decision corresponds to the ‘winning’ population, which here is ‘*D*’.

In this model, decision depends on the cumulative number of spikes, *n*^*U*^(*t*) and *n*^*D*^(*t*), emitted by populations *U* and *D* up to time *t* in a trial. A decision is made at time *t*\*, at which the absolute value of the difference in the numbers of spikes, |*Δn*(*t*\*)| = |*n*^*U*^(*t*) − *n*^*D*^(*t*)|, reaches a given threshold, *θ*, for the first time. The decision is ‘Up’ if *Δn*(*t*\*) = *θ*, whereas it is ‘Down’ if *Δn*(*t*\*) = −*θ* (Fig. 3b).

The psychometric curve of an example network is depicted in Fig. 4a (center; black). Because of the dependence of the firing rate distributions on *ΔL*, the larger *ΔL* the more likely it is that the network would choose ‘Up’. However, the outcome of this decision process is not deterministic. Because spiking is stochastic, *Δn*(*t*) occasionally reaches the threshold that is incongruent with the stimulus, resulting in an error. More generally, because of this stochasticity, the psychometric curve is a smooth sigmoidal function of *ΔL* rather than a step function. Note that in the black psychometric curve of Fig. 4a (center), the network’s perceptual decision in the “impossible trials” is approximately at chance level. Thus, this particular network does not exhibit a substantial ICB.

**Figure 4:**
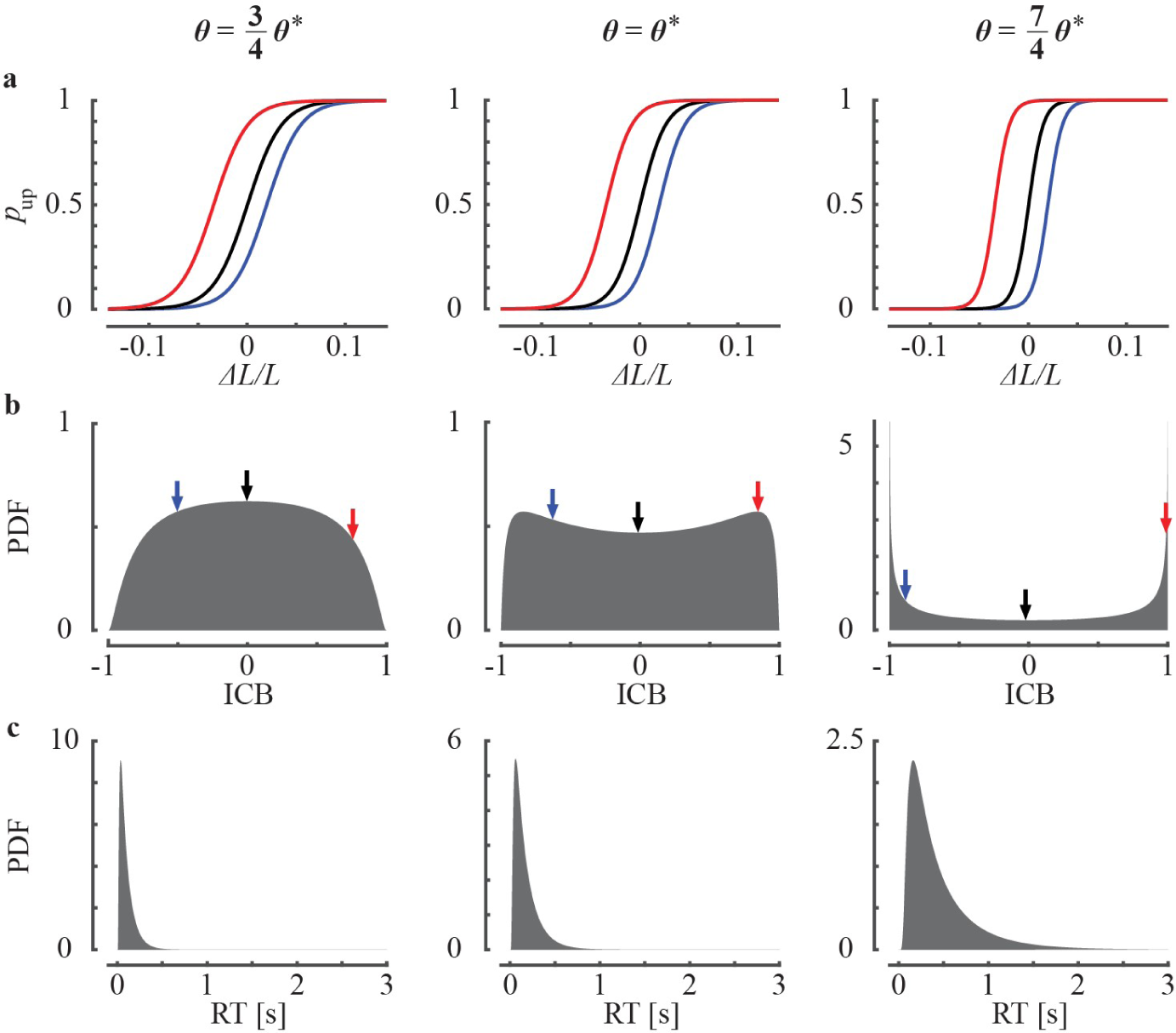
ICBs in the Poisson network model. Left, center and right correspond to network behaviors with low, intermediate and high thresholds (See Materials and Methods). **a**, The psychometric curves of three networks. Each color corresponds to a different network and the same color in different panels corresponds to the psychometric curves of the same network with different thresholds (Eq. 2 in Materials and Methods). **b**, ICB distributions (Eq. 3 in Materials and Methods). The ICBs of the three networks in (a) are denoted in the histogram by arrows of corresponding colors. c, Distribution of reaction times (RTs) (Eq. 4 in Materials and Methods). Number of neurons in the network is *N* = 200,000, 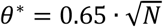.

The black psychometric curve in Fig. 4a (center) was obtained for a particular realization of the network. The red and blue lines in Fig. 4a (center) depict the psychometric curves of two other realizations of the network. Despite the fact that the three networks were constructed in the same way, i.e., by randomly drawing the firing rates of the neurons from the same distributions, the red and blue curves are horizontally shifted relative to the black psychometric curve. Thus, in contrast to the “black” network, the “red” and “blue” networks exhibit ICBs in favor and against responding ‘Up’. The distribution of the ICBs across networks is depicted in Fig. 4b (center). It demonstrates that a wide distribution of ICBs naturally emerges in this decision network model.

It is possible to mathematically prove that for a large network, the behavior of this Poisson network model is equivalent to that of a DDM with a biased drift (Materials and Methods). The accumulation of the difference in the spike counts can be mapped to the accumulation of noisy evidence in the DDM; trial-to-trial variability results from the stochasticity in the neuronal firing; the drift bias stems from the heterogeneity in the firing rates in the two populations. In what follows, we provide an intuitive explanation for the emergence of ICBs in the Poisson network model.

A wide distribution of ICBs in a network consisting of a small number of neurons is easy to understand. Let us consider the impossible trials in a network composed of only two Poisson neurons (Eq. 1 in Materials and Methods), each representing one choice (‘Up’ or ‘Down’). The firing rates of the two neurons are independently drawn from the same lognormal distribution. However, the actual firing rates of these two neurons will, in general, differ in any given network. In some realizations of the network the firing rate of the ‘*U*’ neuron will be higher than that of the ‘*D*’ neuron, whereas in others, it will be lower. Choice is determined by the first threshold-reaching of the accumulated difference in the number of spikes fired by the neurons. It will more often be congruent with the neuron whose firing rate is higher. However, because the firing of spikes in the model is stochastic, the decision in a minority of trials decision will be incongruent with that neuron. This argument implies that this two-neuron network exhibits an ICB, which results from the interplay between the Poisson noise and the heterogeneity in the firing rates of the two neurons. The spiking stochasticity decreases the bias, whereas the firing-rate heterogeneity increases it.

It is thus clear why ICB is natural in such small decision-making networks. However, it is not immediately clear why ICBs are observed in Fig. 4a, in which the number of neurons in the network is large (*N* = 200,000). In this case, the difference between the population averaged firing rates of the U and D neurons is vanishingly small. This is ^because in large networks, this difference is of the order of 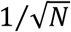, where *N* is the number of neurons. Since the competition between the two populations, which underlies the decision making, is a macroscopic process, one may expect that only average properties of the two populations would affect its outcome. Thus, heterogeneity in the firing rates should not play a significant role in the decision process in large networks. One should note, however, that the sensitivity of the decision-making process to the firing rate heterogeneity increases in proportion to 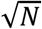. This is because^ the spike times are independent between neurons and thus the fluctuations in spike count decrease with *N*. As a result, the larger the network, the more sensitive it is to the firing rate heterogeneity. Because the sensitivity to the heterogeneity increases in proportion to 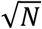 while the heterogeneity itself is proportional to 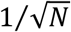, the effect of the heterogeneity in firing rates on behavior is independent of *N* (in the limit of large *N*). Thus, even if *N* is very large, the distribution of ICBs is wide (Materials and Methods).

Unlike network size, the decision threshold has a large effect on the magnitude of the ICB. This is depicted in Figs. 4a, where the psychometric curves of three networks, only differing in the value of the decision threshold, are plotted. The larger the threshold, the steeper is the psychometric curves. This is because the time it takes the network to reach a decision increases with the threshold (Fig. 4c). Thus, a larger threshold results in the integration of spikes over longer durations before a decision is made. Therefore, the decision outcome is less sensitive to the Poisson noise. On the other hand, the network heterogeneity is independent of decision time. Because the magnitude of the ICB is determined by the interplay of the Poisson noise and networks heterogeneity, the larger the threshold is, the broader will be the distribution of ICBs (Fig. 4b; see also eq. (5) and Fig. S4c-d).

### ICB in the recurrent spiking network

Our analysis of the Poisson model suggests that decision making networks exhibit ICBs if (1) the constituting neurons fire irregularly (e.g., Poisson), (2) the neuronal firing rates are heterogeneous (e.g., log-normally distributed) and (3) decision is based on competition (e.g., threshold crossing of the difference in spike counts). In the Poisson model, these three ingredients are introduced ad hoc. Here we investigate a spiking network model in which these features are emergent properties of the network dynamics.

This model builds on previous studies that have shown that recurrent networks of excitatory and inhibitory neurons connected by numerous and strong synapses readily operate in a regime in which excitation is dynamically balanced by inhibition^26^. Two hallmarks of this regime are (1) Poisson-like temporal variability of spike timing (Fig. 5b and S5) and (2) approximately log-normally distribution of firing rates (Fig. 5c). These features emerge from the intrinsic deterministic dynamics of the network (even when the neurons are identical and receive the same external input)^27,28^. Similar to previous models of decision making^29,30^, competition between the alternative actions in our model is mediated by inhibition.

**Figure 5:**
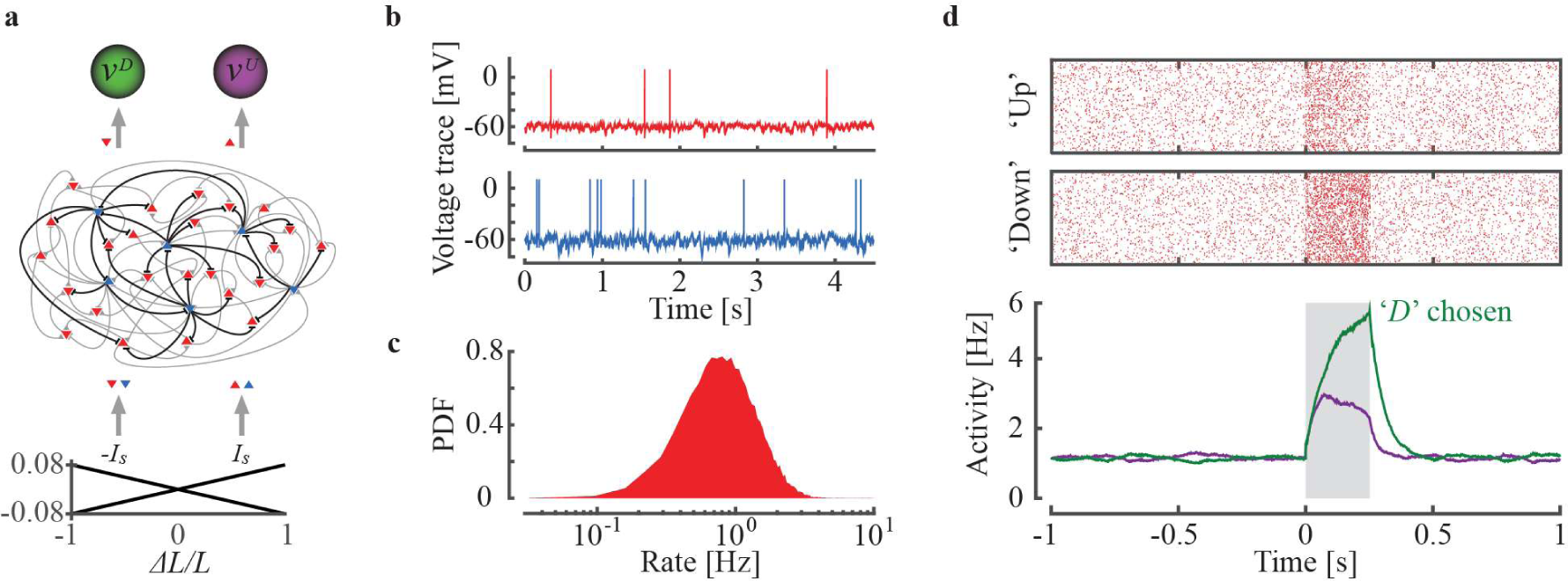
The recurrent spiking network model. **a**, Schematic illustration of the network architecture. The network consists of recurrently connected excitatory (red) and inhibitory (blue) LIF neurons receiving stimulus-selective feed-forward input (bottom). Direction of triangle indicates the selectivity. **b**, Spontaneous activity of example excitatory (red) and inhibitory (blue) neurons in the network. **c**, Distribution of the spontaneous firing rates of the 32,000 excitatory neurons in the network. **d**, Raster plot (10% of the excitatory neurons, top) and the average firing rates (bottom) of the excitatory ‘*U*’ (purple) and ‘*D*’ (green) neurons in response to a stimulus (gray region). The decision is made when the relative difference between the firing rates of the excitatory neurons of the two populations crosses the decision threshold (after which the feed-forward input ceases).

Our model consists of 32,000 excitatory and 8,000 inhibitory Leaky Integrate and Fire (LIF) neurons (Fig. 5a; see Materials and Methods). All neurons receive a feedforward input, which is selective to the stimulus. For half of the neurons (‘*U*’ neurons), this input linearly increases with *ΔL*, whereas for the other half, (‘*D*’ neurons), it is a decreasing function of *ΔL* (Fig. 5a, bottom). When the two segments are of equal length (impossible trials), the ‘*U*’ and ‘*D*’ neurons receive the same feedforward input. All neurons are recurrently connected by strong synapses in a random and non-specific manner, i.e., independent of the selectivity properties of the pre- and post-synaptic neurons. The competition between the ‘*U*’ and the ‘*D*’ neurons is mediated by an additional set of inhibitory connections, which are functionally specific, less numerous but stronger than the unspecific ones (Fig. 5a center; black, specific; gray, non-specific). To investigate the dynamics of this model we performed numerical simulations (See Materials and Methods).

Figure 5d depicts the spike times of 3,200 excitatory neurons in one ‘impossible’ trial. Before the stimulus is presented (*t* < 0), the activities of the ‘*U*’ and ‘*D*’ neurons are similar (Fig. 5d). In response to the sensory stimulus (*t* = 0), the neurons in both populations increase their firing rates. Because of the competition induced by the specific inhibitory connectivity, population ‘*D*’ inhibits population ‘*U*’ and as a result population ‘*U*’ disinhibits the excitatory neurons in population ‘*D*’. In our model, the decision occurs once the relative difference in the average firing rates of the excitatory neurons of the ‘*U*’ and the ‘*D*’ populations exceeds a threshold. After the decision is made, the feedforward stimulus-dependent input ceases and the network activity reverts to its baseline levels (Fig. 5d; see also Materials and Methods).

Fig. 6a (center, black) depicts the psychometric curve of the specific network simulated in Fig. 5d. When the magnitude of *ΔL* is large, the perceptual decision of the network is almost always correct; as *ΔL* decreases, the error rate increases. Considering the “impossible trials” (*ΔL* = 0), the network’s perceptual decision is approximately at chance level. However, different realizations of the connectivity matrix yield psychometric curves, which are laterally shifted (red and blue curves in Fig. 6a). In contrast to the “black” network, the “red” and “blue” networks exhibit substantial ICBs. To estimate the distribution of ICBs in our recurrent network model, we simulated 200 networks, which only differed in their realizations of the connectivity matrix. We computed the ICB of each network from its choices in 500 “impossible” trials. The center panel in Fig. 6b depicts the distribution of these ICBs across the 200 networks. It is significantly wider than expected by chance (p<10^-6^, one-sided bootstrap test, fair Bernoulli process).

**Figure 6:**
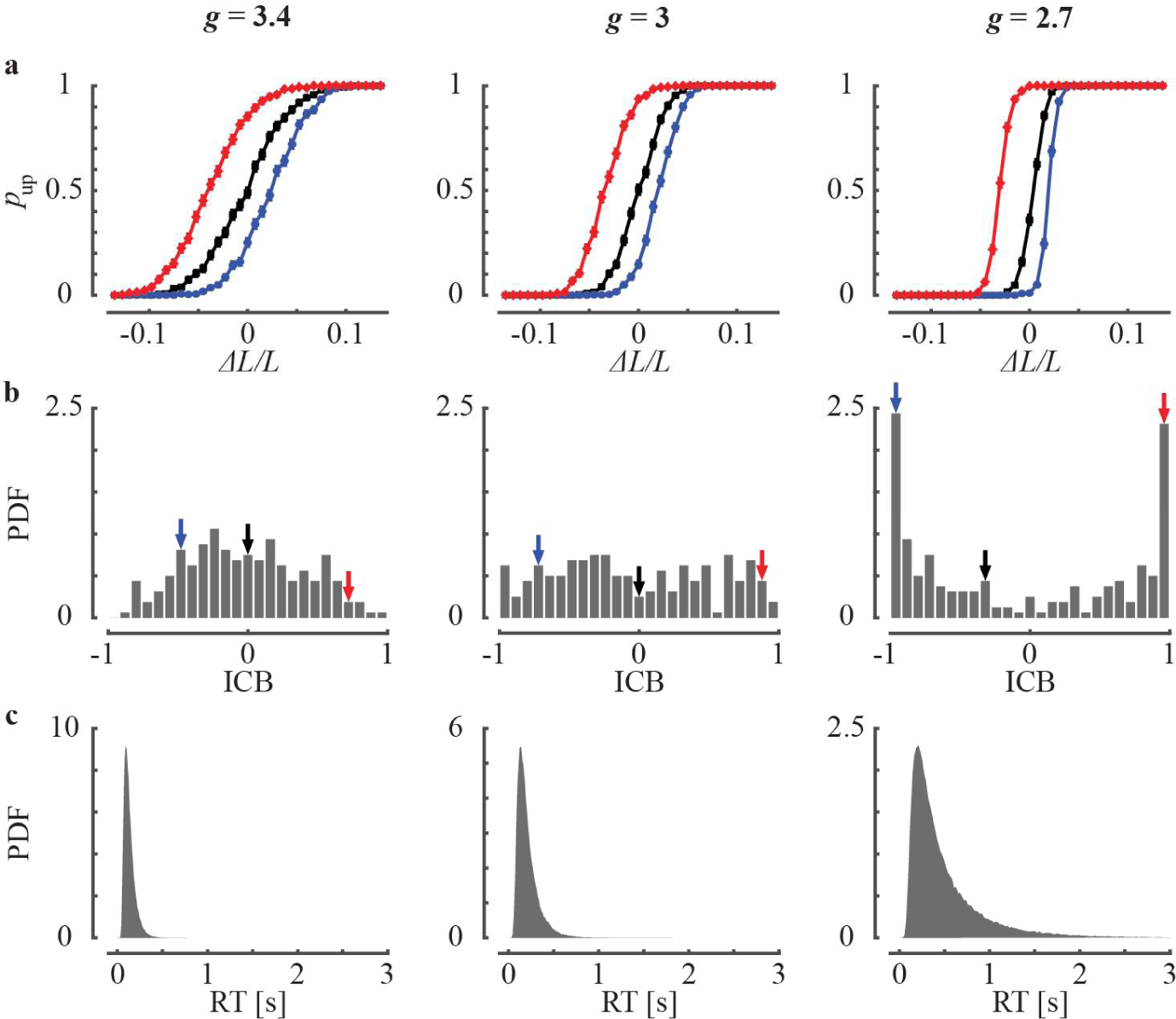
ICBs in the recurrent spiking network model. The strength of the selective inhibition is: Left, *g* = 3.4; Center, *g* = 3.0; Right, *g* = 2.7. **a**, The psychometric curves. Each color corresponds to a different network and the same color in different panels corresponds to the psychometric curves of the same network but with a different *g*. Each point is an average over 500 trials. Error bars correspond to SEM. **b**, Distribution of ICBs for 200 networks. Arrows correspond to the specific psychometric curves in (a) (same color coded). **c**, Distribution of reaction times (RTs).

The level of competition in our model is determined by the strength of the functionally specific inhibition, *g* (see Materials and Methods). Figure 6b depicts the distribution of ICBs for three values of *g*. As *g* increases, the width of the distribution decreases and its shape changes from concave to convex. The distribution of decision times also varies with *g*. The larger *g* the faster is the average decision time (Fig. 6c). When the recurrent network model is analyzed in the framework of the ‘drift bias’ DDM, decreasing the specific inhibition *g* manifests primarily as an increase in the decision threshold (Fig. S6).

It is instructive to analyze the recurrent network model in the framework of the DDM, as we did for the experimental data. Fitting the DDM to the recurrent network simulations, we found that a ‘drift bias’ DDM better explains the network dynamics than ‘IC bias’ DDM (Fig. S7a). Moreover, when the relative contribution of the drift bias and IC bias are tested in the ‘IC+drift bias’ DDM, the contribution of drift bias dominates the emergent ICBs (Fig. S7b). These results are similar to those observed in the behavioral data (Figs. 1d and 2d).

### The conditional bias function

Responses in decision-making tasks can also be analyzed using the *conditional bias function* (CBF). This function quantifies the relationship between bias magnitude and reaction time within the responses of the decision-maker^19^. In the ‘bias drift’ DDM, the distributions of decision times for congruent and incongruent choices are equal^17,31–33^. Therefore, bias is independent of the decision time. By contrast, in the ‘IC bias’ DDM, the bias decreases with decision time^19,20^.

We applied the CBF analysis to the impossible trials in our tasks. Figure 7a depicts the CBF of the responses in the spiking network model, averaged over the 200 networks of Fig. 6. We found that the magnitude of the choice bias decreases with reaction time. The larger *g*, the more negative is the slope of the CBF (Fig. 7a, inset). Because a dependence of the bias on reaction time is expected from asymmetric initial conditions in the DDM, we compared the CBF to that predicted from the fitted ‘IC+drift bias’ DDM. We found that the dependence of the bias on the reaction time in the recurrent model is not well-captured by the fitted ‘IC+drift bias’ DDM (Fig. 7b).

**Figure 7:**
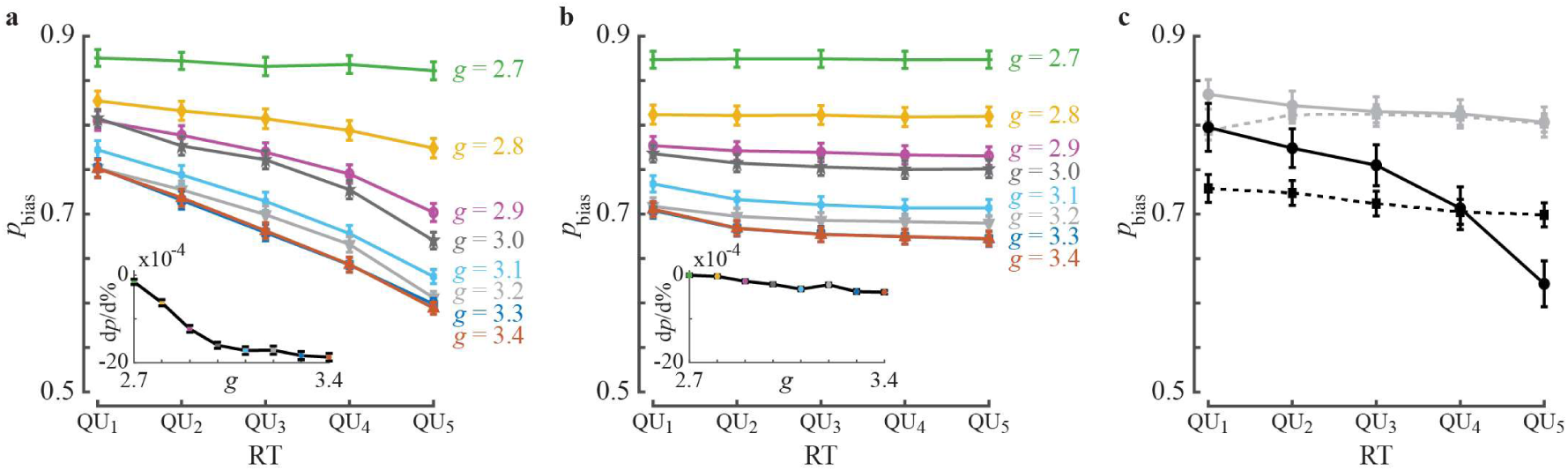
Conditional bias functions. **a**, The recurrent network model. We numerically simulated the recurrent network model in the impossible trials. Responses of each network were divided into 5 quantiles (quintiles) according to the reaction time (RTs). Within each quintile, the fraction of choices congruent with the overall bias of the network, *p*_n717_, was computed and averaged over the different networks (*n* = 200 networks, 500 trials per network). This analysis was performed independently for different values of *g* (different colors; data for *g* = 2.7, 3.0, 3.4 are the same as in Fig. 6). Inset, the slopes of the different curves, defined as the change in *p*_bias_ per percentile of reaction time, *dp*_bias_/*d*%. **b**, ‘IC+drift bias’ DDM fits to the recurrent networks. Same analysis as in (a) was performed on the fitted ‘IC+drift bias’ DDM to the networks’ responses using 10,000 simulated responses for each network. c, Responses of human participants. Solid lines, the conditional bias function for the impossible trials in the bisection task (black) and the motor task (gray). Dashed lines, the corresponding conditional bias functions for the fitted ‘IC+drift bias’ DDMs, based on 2,000 simulated responses for each decision-maker. Error bars are SEM.

We hypothesize that the discrepancy between the CBF of the recurrent network model and that of the corresponding ‘IC+drift bias’ DDM as resulting from a qualitative difference in the decision process in the DDM and the recurrent network. Competition in the recurrent network but not in the DDM results in an effective positive feedback. As a result, evidence accumulated in the beginning of the decision process has a larger effect on the responses than the evidence accumulated later in that process.

This discrepancy prompted us to compute the CBF in our behavioral data. As shown in Fig. 7c, in both the bisection and motor tasks the dependence of the bias on the reaction time is stronger than expected from the fitted ‘IC+drift bias’ DDM. These results indicate that the recurrent network model captures additional features of the behavior, beyond the ‘IC+drift bias’ DDM.

## Discussion

We experimentally investigated human ICBs in a discrimination task and in a motor task. We analyzed the behavior of the participants in the framework of the DDM. We found that in this framework, idiosyncratic biases in the drift rate account for these ICBs. We proved mathematically in a particular model that ICBs due to idiosyncratic drift biases naturally emerge in a network characterized by (1) irregular firing of the neurons, (2) heterogeneity of their firing rates and (3) competition. Finally, we constructed a recurrent network model of spiking neurons, in which these three features are the result of the deterministic dynamics. We numerically simulated this network and demonstrated that it exhibits ICBs, whose features are similar to those observed experimentally. Taken together, our results show that ICBs naturally emerge from the intrinsic dynamics of decision-making neural circuits.

### Bias in the DDM

In the framework of the DDM, choice bias can emerge either from a drift-bias in the decision variable or from asymmetry in its initial condition^19–22^. There is an active area of research that maps different factors affecting choice bias to these two mechanisms. In perceptual tasks, stimuli affect responses via the drift rate^13^. Other factors such as decision cutoff^19,21^ and the bias induced by the action taken in the previous trial^34^ have both been primarily associated with biases in the drift rate. Arousal levels also affect the magnitude of choice biases through the drift rate^35^. By contrast, asymmetries in the prior distribution of stimuli or in the reward schedule predominantly manifests as an asymmetric initial condition^19,21,22,36^ (but see^37^). Heterogeneity among the participants along any of these factors is expected to result in ICBs. Our modeling work predicts the existence of additional, irreducible, ICBs. These ICBs cannot be explained by the experimental context and manifest as drift rate idiosyncrasies in the DDM. Our experimental work reports such ICBs in a discrimination task and a motor task.

### The temporal-scale of stochasticity

In the two models that we have investigated, ICBs emerge from the interplay of two sources of stochasticity: (1) Stochasticity in the timing of action potentials; (2) Heterogeneity in the neuronal firing rates. Stochasticity in the timing of action potentials differs between trials and therefore we can refer to it as fast noise. By contrast, the second source of stochasticity is the same in all trials and therefore, we can refer to it as frozen noise. Cortical dynamics exhibits additional time-scales^38^. Incorporating additional time-scales to the models will not qualitatively affect the results, as long as the contributions of these additional sources of stochasticity are of the order of 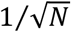 (where *N* is the number of neurons in the network).

To identify the potential contribution of stochasticity at minutes’ time-scale, we tested whether ICBs differed between the first and second halves of our experiments. We did not observe changes in the ICBs in this time-scale that are statistically significant (vertical bisection + motor: two-sided permutation test identified significant, differences in only 17/300 of the pairs, *p* < 0.05, not corrected for multiple comparisons). It will be interesting to quantify the dynamics of ICBs over longer time-scales.

### The effect of correlations

In the Poisson model, spikes are uncorrelated in time and between neurons. As a result, for sufficiently large networks, the magnitudes of both fast and frozen sources of stochasticity decrease as 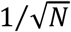. Their ratio and hence the distribution of ICBs become independent of *N* for sufficiently large networks. The two sources of stochasticity also satisfy these scalings in the recurrent network model. This is because the network operates in the balanced regime^26,39^. Noise correlations in the spike count of the neurons are therefore very weak^40–42^ and the firing rates are widely distributed and are uncorrelated between neurons^27^. In network exhibiting correlations in the neuronal activity^41,42^, averaging over neurons may not decrease the fast noise and the heterogeneities as 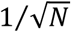. If the dependence of the two on *N* is very different, one source of stochasticity could dominate, resulting in deterministic or unbiased choices.

### Alternative interpretation of the observed ICBs

Idiosyncratic postures can affect visual stimuli or the effort associated with motor responses. One cannot rule out the possibility that such idiosyncrasies contributed to the ICBs in the bisection task, which was performed online. In the motor task, by contrast, all participants were dextral and the positions of the chair, screen and mouse pad were kept the same for all participants. Nevertheless, we cannot exclude the possibility that differences in participants’ anatomy, e.g., their arm length, contributed to the ICBs.

Reinforcers can affect choice preferences^43–45^. The specific history of stimuli also influences preferences in perceptual tasks^9,46^. Along these lines, it is natural to attribute ICBs to the specific histories of the participants during the experiment. We therefore designed our tasks to minimize operant and sequential effects. Nevertheless, we cannot exclude the possibility that the observed ICBs are the result of operant or sequential effects which occurred before the experiment. For example, considering the impossible trials in our bisection task, participants may prefer to press the Down key because they are accustomed to pressing taskbar icons that are located at the bottom of their computer monitor. Other participants may prefer the Up key because they are used to a taskbar located at the top of the screen. In such a view, ICBs in the vertical bisection task can be attributed to idiosyncratic histories of computer usage prior to the experiment.

All of the above effects can contribute to ICBs. However, we showed that these and similar explanations are not necessary. All that is required for ICBs are minute differences between the populations encoding the two alternatives. Such differences are almost inevitable in any 2-alternative task, in which the two alternatives are represented by different populations of neurons.

Substantial ICBs were observed in genetically-identical flies that were reared in the same environment^47^. The results of that study suggest that biases can emerge from effects that are unpredictable from genetic, environmental or anatomical variables. This is in line with our study that showed that the random differences in the fine structure of connectivity between the neuronal populations involved in decision-making are sufficient to account for the ICBs. In conclusion, the occurrence of ICBs in a cortical-based decision task is thus almost inevitable. It would be therefore surprising to find a decision task that is devoid of the ICBs, unless they are actively suppressed, e.g., by penalizing them.

## Materials and Methods

### The perceptual discrimination task

The study was approved by the Hebrew University Committee for the Use of Human Subjects in Research. Recruitment was based on the online labor market Amazon Mechanical Turk^48^. Data were collected from 100 participants (51 males, 49 females; 91 dextrals, 7 sinistral, 2 ambidextrous; mean age = 39 years, min = 22 years, max = 71 years). All participants were Mechanical Turk’s Masters, located in the United States of America. All participants reported normal or corrected to normal vision and no history of neurological disorders. The experiment was described as an academic survey of visual acuity. A base monetary compensation was given to all applied participants for the participation. In order to encourage good performance throughout the experiment, an additional bonus fee was given for every correct response and another bonus was guaranteed to 10% of participants with highest scores.

#### Procedure

Participants were instructed to indicate the offset direction of the transecting line, out of two alternative responses. Possible responses were either ‘Left’ or ‘Right’, for the horizontal discrimination task, or ‘Up’ or ‘Down’, for the vertical discrimination task. Participants were asked to answer as quickly and accurately as possible.

In each trial, a 200 pixel-long white line, transected by a perpendicular 20 pixel-long white line was presented on a black screen (Fig. 1a, inset). The stimuli were limited to a 400-pixel X 400-pixel square at the center of the screen. Window resolution was verified for each participant individually, to make sure that it did not exceed the centric box in which all stimuli were presented. The horizontal location of all vertical bisection lines and the vertical location of all horizontal bisection lines were centered. After 1 sec, the stimulus was replaced by a decision screen composed of two arrows, appearing in opposite sides of the screen, and a middle 4-squares submit button. The participants indicated their decision by moving the initially centered cursor to one of the arrows, pressing it, and finalizing their decision by pressing the ‘submit’ button. No feedback was given regarding the correct response. The participants were, however, informed about the accumulated bonus fee every 30 trials.

The experiment consisted of 240 trials, 120 horizontal and 120 vertical. Trials were ordered in 80 alternating blocks of 3 horizontal and 3 vertical transected lines. Unbeknown to the participants, there were 20 impossible horizontal and 20 impossible vertical trials (⅙ of the trials). To minimize sequential effects in the impossible vertical bisection trials, each impossible vertical bisection trial was preceded by three horizontal bisection trials. The order of the trials was pseudorandom but identical for all participants. For the possible trials, the deviation from the veridical midpoint was uniformly distributed between 5 and 10 pixels (|∆*L*|/*L* between 0.05 and 0.1, where *ΔL*/*L* ≡ (*L*^*U*^ − *L*^*D*^)−(*L*^*U*^ + *LD*) and *L*^*U*^ and *L*^*D*^ denote the lengths of the Up and Down segments of the vertical line). with an equal number of offsets in each direction. Because it is well established that in the horizontal bisection task participants exhibit a global bias (attributed to pseudoneglect^49^), we focused on the vertical bisection trial in quantifying ICBs and performance. Mean performance in the possible vertical trials was 96.4% ± 4.6% (standard deviation), range 71% − 100%. No participants were excluded from the analysis. In the DDM analysis we excluded trials, in which the reaction time was longer than 3 sec. This excluded 1% of the vertical bisection trials.

To verify that the participants understood the instructions, they were required prior to the experiment to successfully complete a horizontal-bisection practice session and a vertical-bisection practice session. A session consisted of blocks of 4 easy trials (|*∆L*|/*L* = 0.2) with feedback and balanced polarity of ∆*L*. The main experiment started after the participant completed one horizontal and one vertical block successfully. Responses in this practice session were not included in the analysis.

### The motor task

The study was approved by the Hebrew University Committee for the Use of Human Subjects in Research. The experiment was described as an academic survey testing speed of motion. Data were collected from 20 participants (13 males, 7 females; all dextrals; mean age = 25 years, min = 19 years, max = 41 years) who were recruited using on-campus advertising. All participants reported normal or corrected to normal vision and no history of neurological disorders.

#### Procedure

In each trial, a pair of dots, equally distant from a central black disk, were presented on a background of a larger white disk (Figs. 2a and S3a). Participants were instructed to drag as quickly as possible the two dots into the black disk using the mouse cursor. Each trial started with a forced delay period of 0.75 sec. Then, the mouse cursor appeared in the center of the disc. The participant used the mouse to move the cursor to one of the dots. She then dragged the chosen dot to the central black disk by pressing the mouse and moving it. If accurate, a release of the dot on the central black disk resulted in a 1.1 sec “swallowing” of the dot animation, indicating a successful drag. The dragging time (measured from the time of clicking on the dot to the time of its release) appeared on the screen. It disappeared after a forced delay of 1.1 sec and the cursor reappeared in the center of the disk. The participant processed the second dot in the same way as the first dot. We used 10 different pairs of dots, each presented 20 times. Each pair of dots was of equal distance from the center of the black disk, but of a different color and a different angular location (Fig. S3b). The order of presentation was pseudorandom such that in every consecutive group of 10 trials all pairs appeared. Decision time in a trial is defined as the time elapsed from cursor appearance to the beginning of the dragging of the first dot. The positions of the chair, screen and mouse pad were fixed and identical for all participants in order to minimize heterogeneity between participants.

A base monetary compensation was given to all participants for their participation. An additional bonus fee was given based on dragging times in order to encourage good performance throughout the experiment.

In the DDM analysis we excluded trials, in which the reaction time was longer than 3 sec. This excluded 2% of the motor trials.

### The Poisson model

We consider two populations of neurons, denoted by *‘U’* and *‘D’*, representing choice ‘Up’ and ‘Down’ (Fig. 3a). Each population consists of *N*/2 independent Poisson neurons. The stimulus-dependent feedforward inputs to neuron *i* (*i* ∈ {1, …, *N*/2}) in population *α* (*α* ∈ {*U*, *D*}) is given by: 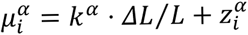, where *k*^*U*^ = −*k*^*D*^ = *k* is a parameter and 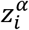 is stimulus- and trial-independent, independently drawn (once) from a zero-mean Gaussian distribution with variance 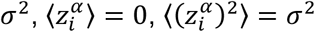, where 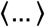 denotes average. The firing rate 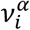, different for each neuron, is

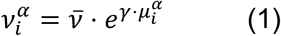

where 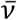 is a baseline firing rate and *γ* is the gain^27^. Due to the exponential transfer function and the normal distribution of inputs, the firing rates are log-normally distributed. In each trial, the cumulative number of spikes, *n*^*U*^(*t*) and *n*^*D*^(*t*), emitted by populations ‘*U*’ and ‘*D*’ up to time *t* in a trial is counted (Fig. 3a). A decision is made when (*t*\*) the absolute value of the difference in the numbers of spikes, |*Δn*(*t*\*)| = |*n*^*U*^(*t*) − *n*^*D*^(*t*)|, reaches a given threshold 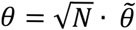, where 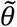 is a parameter.

For *N* ≫ 1 and neglecting the threshold effect, the difference in spike count at time *t* is given by *Δn*(*t*)~*𝓝*(*Δv* ∙ *t*, Σ*v* ∙ *t*), where 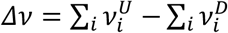 and 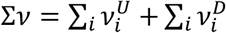. Because *N* ≫ 1, both *Δv* and Σ*v* are normally distributed:

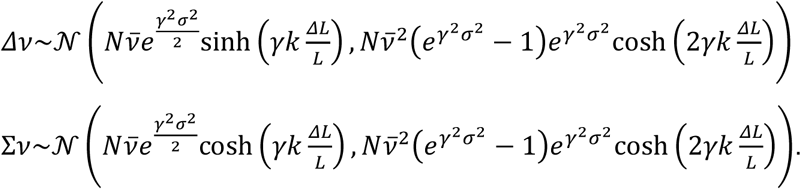

Note that *Δn* and *Δv* are different stochastic processes: the stochasticity of *Δn* stems from trial-by-trial variability, conditioned on the firing rates of the neurons. By contrast, the stochasticity of *Δv* reflects heterogeneity in these firings rates across different realizations of the decision-making network.

The standard deviation of the distribution of Σ*v* is of 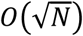, whereas its mean is *O*(*N*) even when *ΔL* → 0. Therefore, in the limit *N* ≫ 1, 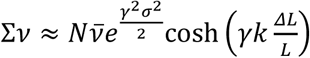. By contrast, in the regime in which 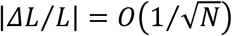, the mean and standard deviations of the distribution of *Δv* are comparable, both are 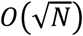.

The probability of an ‘Up’ decision is obtained by solving a first-passage problem, yielding

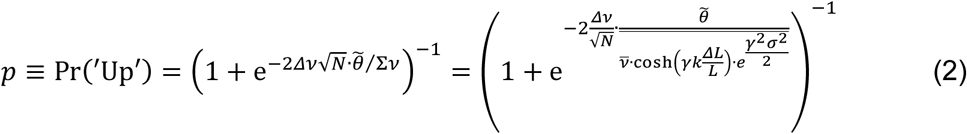

Substituting the dependence of *Δv* on *ΔL* in Eq. (2) yields the psychometric curve. In particular, when the two networks are symmetric, 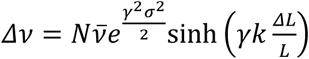, and 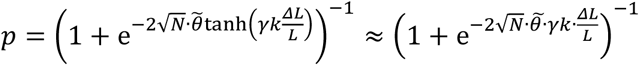

More generally, when the two networks are only drawn from the same distribution, the psychometric curve will be horizontally shifted relative to the identical networks case.

To compute the distribution of ICBs, we consider the case in which the external input is symmetric, *ΔL* = 0 and thus 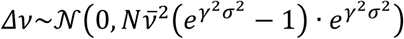. After a change of variables,

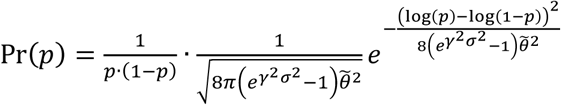

Because ICB = 2*p* − 1,

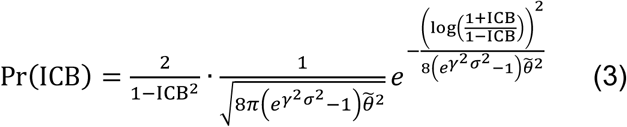

The corresponding distribution of decision times is computed by averaging the drift-conditioned distribution of first-passage times over the distribution of *Δv*, yielding^17,31,32^:

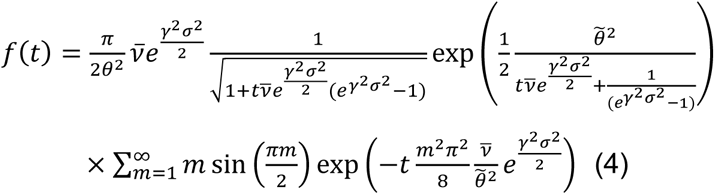

Two points are worthwhile noting:

1. The neuronal gain parameter *γ* affects Pr(*p*) through the term 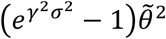. This implies that increasing the gain is effectively equivalent to increasing the threshold parameter 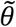, and thus is likely to broaden the distribution of ICBs.
2. The assumption of a lognormal distribution of firing rates is not essential to our analysis. For a general distribution of firing rates, Eq. (3) becomes

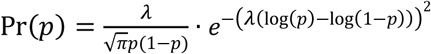

and

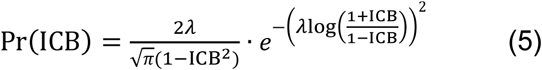

where 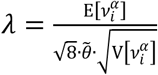 and 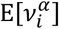 and 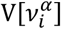 are the mean and variance of the distribution of the firing rates in the impossible trials.

The parameters used in all simulations are 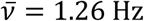, *γ* = 1, *k* = 0.133 and *σ*^2^ = 1. For *ΔL* = 0, the average and standard deviation firing rate are 2.1 Hz and 2.7 Hz. These numbers are compatible with experimental data in the cortex^25,50^. The dependence of the width of the ICB distributions on the model parameters are depicted in Fig. S4.

### The spiking network model

The model is a recurrent network of *N* leaky-integrate-and-fire (LIF) neurons, *N*^*E*^ = 0.8*N* excitatory and *N*^*I*^ = 0.2*N* inhibitory (the superscript denotes neuron type, excitatory or inhibitory).

*Single neuron dynamics*: The sub-threshold dynamics of the membrane potential, 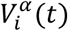, of neuron *i* in population *α* (*i* = 1, …, *N*^*α*^; *α* = *E, I*) follow:

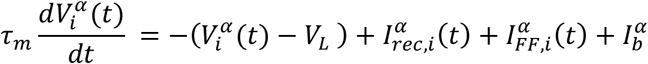

where *τ*_*m*_ is the neuron membrane time constant, *V*_*L*_ is the reversal potential of the leak current. Inputs to the neuron are modeled as currents: 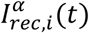 is the recurrent input into neuron (*i*, *α*), due to its interactions with other neurons in the network, 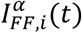 is the feedforward input into that neuron elicited upon presentation of the stimulus, and 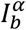 is a background feedforward input, independent of the stimulus, identical for all the neurons and constant in time. These subthreshold dynamics are supplemented by a reset condition: if at 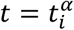 the membrane potential of neuron (*i*, *α*) reaches the threshold, 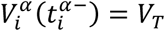, the neuron fires an action potential and its voltage resets to 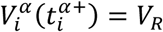.

*The feedforward input*: Each population, excitatory or inhibitory, consists of two types of neurons, namely *U*- and *D*-selective. In the absence of stimulus, the feedforward input 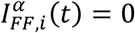 for all the neurons. Upon presentation of a stimulus for which *ΔL* > 0, 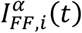 into U-selective neurons is stronger than 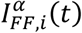 into D-selective neurons. The opposite is true when *ΔL* < 0. Specifically, we take:

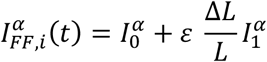

where 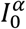 and 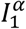 are constants and positive and *ε* characterizes the selectivity of the neuron: *ε* = +1 for ‘*U*’ neurons and *ε* = −1 for ‘*D*’ neurons. We denote the set of U-selective (resp. D-selective) neurons in population *α* = *E, I* by *U*^*α*^ (resp. *D*^*α*^). Neuron 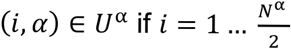 and 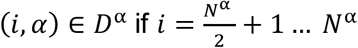.

*The recurrent input*: The connectivity has two components. One is functionally specific and the other is not. The non-specific component is fully random (Erdös-Renyi graph) and does not depend of the selectivity of the pre- and post-synaptic neurons. The corresponding *N*^*α*^ × *N*^*β*^ connectivity matrix, 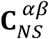, is such that 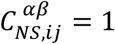 with probability *k*/*N*^*β*^ and 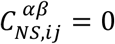 otherwise, where *K* is the average number of non-specific inputs that a neuron receives from neurons in population *β*. The strength of the non-specific connections depends solely on *α*, *β* yielding: 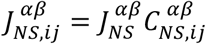 where 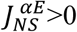 (excitation) and 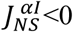 (inhibition).

The competition between the ‘*U*’ and the ‘*D*’ selective neurons is mediated by an additional set of connections. These connections are specific and are much less numerous but stronger than the unspecific ones. The corresponding connectivity matrices, 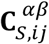, are such that:

1. 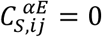 i.e. we assume no specific excitation.
2. 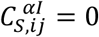 if *i* and *j*; have the same selectivity properties.
3. 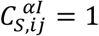 with probability 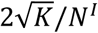 if *i* and *j* have different selectivity properties.

Therefore, each neuron (excitatory as well as inhibitory) receives, on average, 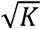 connections from inhibitory neurons whose selectivities are different from its own (compared with, on average, *K* non-selective inhibitory connections).

The strength of the specific connections depends solely on the neurons’ type 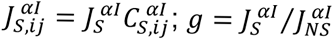.

The total current into neuron (*i*, *α*) due to the recurrent interactions is

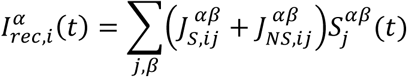

where 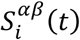 are synaptic variables, which follow the dynamics

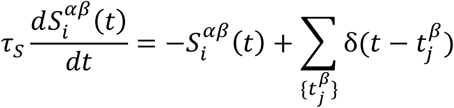

Here, *τ_s_* is the synaptic time constant (assumed to be the same for all synapses) and the sum is over all spikes emitted at times 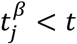.

*Decision-making and decision criterion*: In response to the sensory stimulus, the activities of the *U*-selective and *D*-selective neurons change differently (Fig. 5d). We compute at every time step the population-averaged activity of all the excitatory neurons in the set *a*, (*a*-selective), denoted by *v*_*a*_, *a* ∈ {*U*, *D*}, by convolving the spike times with an exponential filter with a time constant of 50 msec. Decision is based on the ratio: 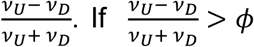, the decision provided by the network is that upper segment is longer than the lower one, whereas for 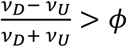 it is the opposite, where *ϕ* > 0 is the decision threshold.

The ability of the network to make a decision depends on the network parameters. In particular, it depends on the parameter *g*, which characterize the strength of the competition between ‘*U*’ and ‘*D*’ neurons, on the value chosen for the threshold *ϕ* as well as on the stimulus parameters, 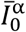 and 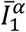.

*Numerical integration*: The dynamics of the model circuit were numerically integrated using the Euler method supplemented with an interpolation estimate of the spike times^51^. In all simulations the integration time step was 0.1 msec. We verified the validity of the results by performing complementary simulations with smaller time steps.

*Model parameters*: A systematic study of the dependence of the network dynamics on the parameters is beyond the scope of the present paper. The parameters used in all the simulations are: *V*_*L*_ = −60mV; *τ*_*m*_ = 10msec; *V*_*T*_ = 10mV; *V*_*R*_ = −60mV; *J*_*EE*_ = 35mV ∙ ms, *J*_*IE*_ = 233.3mV ∙ ms, *J*_*EI*_ = −175mV ∙ ms, *J*_*II*_ = −233.3mV ∙ ms, 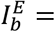,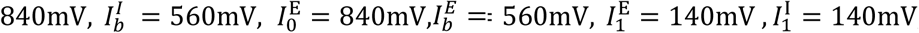, *τ*_*s*_ = 3msec. The total number of neurons and average non-specific connectivity is *N* = 40,000, *K* = 400, *ϕ*= 0.4. The sole parameter we vary is *g*.

The single-neuron parameters and the average number of inputs per neuron are as in^41^. The network size and fraction of inhibitory neurons are as in^28^. The strengths of E→E connections, as well as the unstructured components of I→E and I→I interactions are as in^41^.

The background external inputs as well as the E→I connection strength were chosen to obtain spontaneous and evoked firing rates that are comparable with the experimental data. The decision threshold was chosen so that decision occurs only when the differences in the firing rates of the two populations is comparable to the experimentally observed^15^.

*Relation to other spiking network models*: In previous LIF network models of decision making^30,52^, the recurrent connectivity of the competing populations encoding for the decision outcome is all-to-all and symmetric. In contrast, the connectivity in our network is sparse and random. In these previous studies, irregular firing is due to extrinsic stochastic input. In our model it is generated by the intrinsic recurrent dynamics. Interestingly, it is reported in the Supplementary Methods of^52^ that in their model, “The random connectivity within the sensory circuit and across circuits (i.e. bottom-up and top-down) can cause that the network’s behavioral responses exhibit a bias towards one of the two choice options”. We hypothesize that the mechanism underlying that bias can be understood using our framework.

### DDM analysis

According to the DDM^13–15,17,18,53–55^, noisy evidence in favor of choosing each of the two alternatives is integrated over the course of the trial. The difference of these evidence, a quantity known as the decision variable, is then computed. Mathematically, *dp*/*dt* = *A* + *ξ*, where *x* is the decision variable, *A* is the drift rate, *t* is time within the trial and *ξ* denotes white noise such that E[*ξ*(*t*)] = 0 and E[*ξ*(*t*)*ξ*(*t*′)] = *δ*(*t* − *t*′). In the free-response version of the DDM, which has proven useful for modeling choices even when the stimulus is presented for a fixed duration^56–58^, a decision is made once the decision variable reaches one of two decision thresholds, 0 or *a* > 0. The initial condition is set to *x*(*t* = 0) = *z* ∙ *a*, where 0 < *z* < 1.

We focus on the impossible trials in which ∆*L* = 0. Evidence for *A* ≠ 0 in those trials is interpreted as drift bias; evidence for *z* ≠ 0.5 is interpreted as initial condition bias (IC bias). The two bias mechanisms exhibit distinct patterns of dependence of bias on reaction-times. The effect of ‘IC bias’ is mostly prominent early in the trial and it therefore predicts that faster decisions are more biased than slower ones. By contrast, ‘drift bias’ affects evidence accumulation throughout the trial and the resulting bias affects both fast and slow decisions^19,20^. Therefore, it is possible to dissect the two mechanisms by incorporating the decision times in the analysis.

We fit four different variants of the DDM to the behavioral data and simulations. (1) A baseline DDM with *A* = 0 (because we consider only the impossible trials) and *z* = 0.5. This model has a single parameter, the decision threshold, *a*. To fit the model to the data, a second parameter, which accounts for the component of the reaction-time that is independent of the decision process, *T*_*er*_, is added^56,59,60^. By construction, there are no ICBs in this model and it was used as a baseline for comparison with the other three models. (2) In the ‘IC bias’ DDM *A* = 0, as in the baseline DDM. However, by contrast, *z* is estimated from the data, this is in addition to *a* and *T*_*er*_. (3) In the ‘drift bias’ DDM, we assumed that *z* = 0.5 and estimated *A*, *a* and *T*_*er*_. (4) In the ‘IC+drift bias’ DDM, both *z* and *A* were estimated from the data, in addition to *a* and *T*_*er*_.

#### Hierarchical Bayesian estimation of the DDM parameters

The dataset of the vertical bisection task includes 20 impossible trials performed by 100 participants. The motor task includes 10 datasets (each pair of dots was considered a task and was analyzed separately). Each task was tested on 20 participants, each performing 20 decisions. The recurrent network simulations included 8 datasets, each corresponding to a different level of specific inhibition. Each of these dataset consisted of the responses made by 200 networks, each tested on 500 impossible trials.

We fit each of the four DDM variants to each of the datasets using the HDDM Python toolbox, which allows for the construction of Bayesian hierarchical DDMs^61^. HDDM uses Bayesian Markov-chain Monte Carlo sampling for generating posterior distributions over both subject-level and group-level model parameters, rather than point estimates of only subject-level parameters. To accomplish this, HDDM uses informative prior distributions on the group-level parameters, that constrain the parameters to a plausible range given past experiments^61,62^. As a result, constraining the parameter estimates for individual subjects by group-level inference leads to a better recovery of the true parameters, especially with few trials per subject^61^.

Our analysis required three minor modifications to the code: (1) In the unbiased and ‘IC bias’ DDMs, we posit that *A* = 0, which corresponds to an unbiased drift rate. This is because in the bisection task and the recurrent network simulations we only analyzed the case of ∆*L* = 0. This constraint was lifted in the ‘drift bias’ and ‘IC+drift bias’ DDMs, in which *A* was a free parameter. (2) In the HDDM fitting procedure, the estimation of each of the model parameters is constrained by the informative priors relevant for the group level statistics of the sample’s parameter. Specifically, it is assumed that the mean drift rate is drawn from a normal distribution with a positive mean, *m* = 2, conceivably because behavior is typically studied in possible trials, in which performance is above chance. Because in the impossible trials there is no a-priori reason to assume that one action is more likely than the other, we modified the code such that *m* = 0. Notably, comparable posteriors are obtained also when using *m* = 2 (not shown). (3) The assumptions regarding the width of the prior distribution of initial conditions can constrain the values of the estimated initial conditions in the HDDM fitting procedure, thus limiting the extent to which the initial conditions can capture the ICBs in the DDM. Therefore, we considered a wider prior distribution of initial conditions, by increasing the standard deviation of *σ*_*z*_ in^61^ from 0.05 to 5. Notably, when keeping the standard deviation of *σ*_*z*_ at 0.05, the contribution of initial conditions to the ICBs in the resultant fitted DDMs is even smaller (not shown). As is standard in the HDDM fitting procedure, we allowed 5% of responses to be considered ‘contaminants’^59^, i.e., trials which do not follow the DDM dynamics (e.g. due to attentional lapses).

In order to estimate the posteriors, we ran 12 separate Markov chains with 40,000 samples each. Of those, the first half was discarded as burn-in and to reduce sample autocorrelations, 4/5 of the remaining samples were discarded for thinning. This left 4,000 samples per chain. We computed the 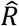 Gelman-Rubin statistic, to assess model convergence by comparing between-chain and within-chain variance of each posterior distribution. For all datasets and all models, the 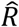 of all group-level posteriors (0.9998-1.01) and that of the observer-level posterior (0.9998-1.035) indicated a proper convergence^63,64^. All chains were concatenated for further analyses, resulting in 48,000 samples per model, from which each posterior was estimated.

#### Model comparison using the DIC

Using the HDDM Python toolbox^61^, we also computed the Deviance Information Criterion (DIC^65^) and used it to compare the different variants of the DDM. The DIC compares models by the goodness of fit, while penalizing for model complexity. The lower the DIC the better the model (see^65^). Because of the nondeterministic nature of hierarchical modeling, we also computed confidence intervals of the *Δ*DIC (DIC of the variant of the DDM relative to the DIC of the baseline, unbiased DDM). For the vertical bisection task and the numerical simulations of the recurrent network, the SEM of the *Δ*DIC was estimated by repeating the fitting procedure and DIC analysis 3 times. For the motor datasets, the *Δ*DIC of each biased DDM variant was obtained separately for each pair of dots, and the SEM was then evaluated over all 10 pairs.

#### Posterior-based simulations

The quality of the DDM models can also be evaluated by comparing the behavior of the decision-maker to the behavior predicted by the estimated posteriors. Specifically, we simulated responses (choices and reaction time) using the posteriors obtained from the HDDM procedure for each dataset separately. For all datasets, the simulated probabilities of choice well-matched the observed ones for the ‘drift bias’ and for the ‘IC+drift bias’ DDM variants but not for the ‘IC bias’ DDM (Fig. S8a). All models provided reasonable fits of the normalized distribution of reaction times to the data (Fig. S8b).

#### Relative contributions of the IC bias and drift bias to the ICBs in the ‘IC+drift bias’ DDM

Here we describe the procedures underlying Figs. 1d, 2d and S7. It is well-known that in the DDM^17,31^

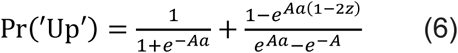

To dissect the relative contributions of the IC and drift biases to the ICBs, we computed the average parameters *A*, *a* and *z* from the estimated posteriors of each observer in the ‘IC+drift bias’ DDM. We then computed the predicted Pr^(^′Up′^)^ in three conditions: all estimated parameters (black Xs), estimated initial conditions *z* ∙ *a* and *A* = 0 (purple squares) and estimated product, *Aa*, of drift with the threshold, while assuming *z* = 0.5 (green circles).

#### Relative contribution of idiosyncratic thresholds to the ICBs

According to Eq. (6), the drift bias *A* and the threshold *a* contribute to the ICB via their product *Aa*. Importantly, while the drift parameter, *A*, can be positive or negative, the threshold parameter, *a*, is strictly positive. Therefore, the direction of the bias is necessarily determined by the drift *A*. Nevertheless, idiosyncrasies in *a* can also contribute to the heterogeneity in the bias between the decision makers. We studied the relative contributions of these two parameters in the framework of the ‘drift bias’ DDM, in which the product *Aa* is the sole contributor to the ICBs. For each decision maker we computed the posterior-averaged values of *a* and *A*. We then used Eq. (6) to predict the ICBs assuming that all decision-makers are characterized by the same (average) threshold or the same (average) drift (|*A*|). In all datasets we found that the contribution to the ICBs of heterogeneity in the thresholds is small relative to that of the drift (Fig. S9).

### The Poisson model is equivalent to the drift bias DDM

Comparing equations (2) and (6), we note that for 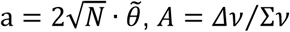 and *z* = 0.5, Eq. (6) is equivalent to equation (2).

### Data availability

The data and the relevant codes can be downloaded from TBD link.

## Acknowledgements

We thank Thomas Boraud, Gianluigi Mongillo, Haim Sompolinsky and Talia Tron for discussions. This work was conducted within the scope of the France-Israel Laboratory of Neuroscience. D. H. thanks the Department of Neurobiology at the Hebrew University for its warm hospitality. This work was supported by the Israel Science Foundation (Y. LO., Grant No. 757/16), the DFG (Y. LO.), the Gatsby Charitable Foundation (Y. LO.), ANR-09-SYSC-002-01 (D. H.) and the France-Israel High Council for Science and Technology (D. H. and Y. LO.).

## Author contributions

L. L., Y. LA., D. H. and Y. LO. conceived and planned the experiments; L. L., R. D., D. H. and Y. LO. developed the models; L. L., D. H. and Y. LO. wrote the manuscript.

