## Supplementary figures for "Idiosyncratic choice bias in decision tasks naturally emerges from intrinsic stochasticity in neuronal network dynamics"

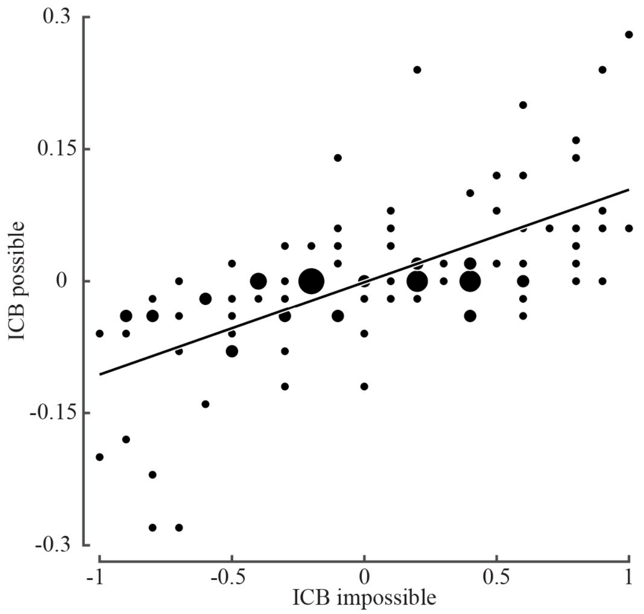


**Figure S1** ICBs in the possible vs. ICBs in impossible trials in the bisection task. For each participant (*n*=100), the fraction of 'Up' responses was computed separately for the possible and impossible trials. Each dot denotes the value of ICB in the possible trials as a function of its value in the impossible trials. Small dots denote single participants. Radii of larger dots denote the number of participants sharing the same values of both possible and impossible ICBs. Black line: best fit orthogonal regression (slope = 0.11). Despite the overall high performance in the possible trials ($96.4\% \pm 4.6\%$), participants exhibited substantial ICBs in the possible trials and these were highly correlated with the ICBs in the impossible trials ($A = \rho = 0.64$, $p<10^{-12}$). Considering only the error trials, in 81% of the error trials (284/351) the error was congruent with the ICB measured in the impossible trials and in only 19% of the error trials (67/351) the error was incongruent with the ICB (correlation of ICB in impossible trials and error trials: $\rho = 0.68$, $p<10^{-10}$; slope of best fit orthogonal regression: 1.88).

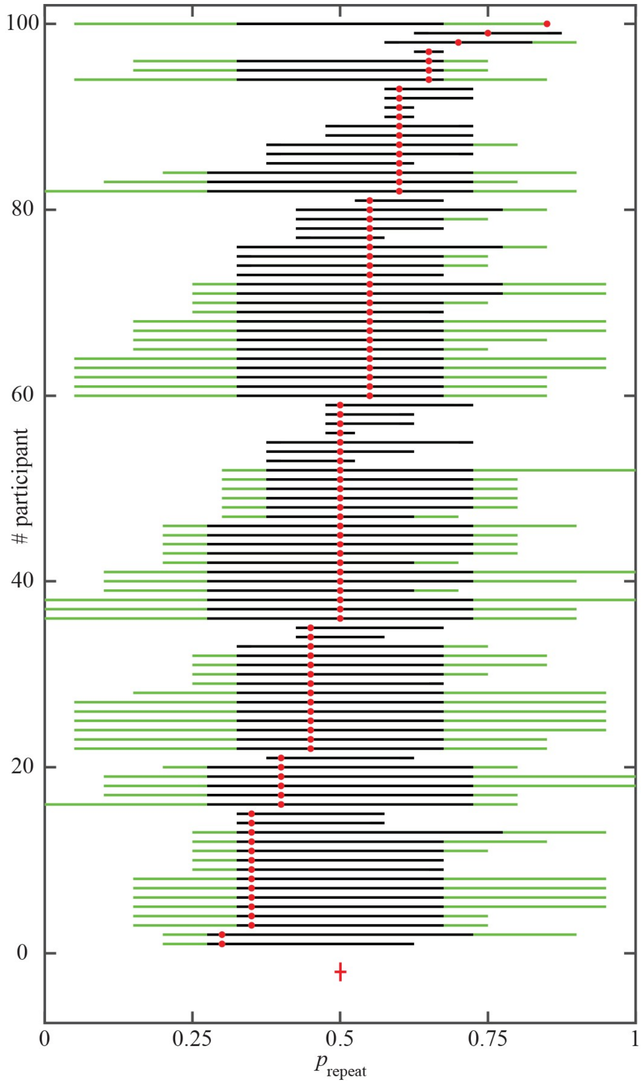


**Figure S2** Lack of evidence for a repetition or alternation tendencies in the bisection task. Each red dot denotes $p_{\text{repeat}}$, the frequency of repeating the decision made in the previous trial. Participants are ordered by $p_{\text{repeat}}$. Black lines denote 90% confidence interval (tested by shuffling the order of decisions, not corrected for the number of participants). Green lines, denote 100% confidence interval (all possible values in the shuffling test). Red error bar: mean $p_{\text{repeat}}$, averaged over all participants ± SEM. Each impossible vertical bisection trial was preceded by three horizontal bisection trials so the previous vertical trial occurred four trials ago. Only one participant (#100) exhibited a statistically significant tendency to repeat.

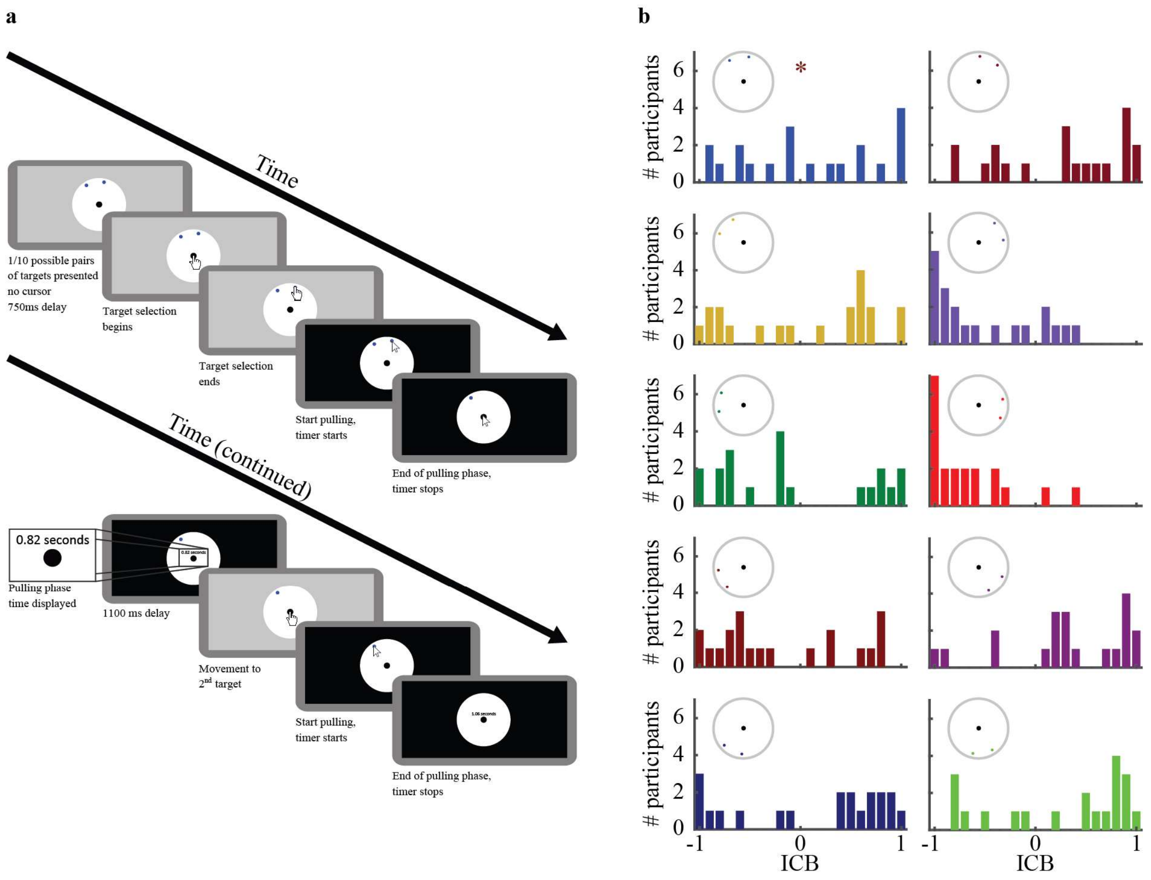


**Figure S3** The motor task. **a**, Detailed description of a single trial in the motor task. A trial began with a presentation of the two dots and a central black disk on a white disk background. After 750 msec, a cursor appeared on the central black disk and the participant used the mouse to place it over one of the dots. Then, the participant clicked on the dot and used the mouse to pull it to the central black disk and released it. An accurate release of the dot resulted in a 1.1 sec "swallowing" of the dot animation. The dragging time (from click to release) was displayed. Participants were instructed to minimize that time. Then, the cursor reappeared on the central black disk and the process repeated. **b**, Locations and colors of all 10 pairs of dots used in the experiment, with the corresponding distributions of choice biases. The pair of dots depicted in Fig. 2a is denoted by an asterisk.

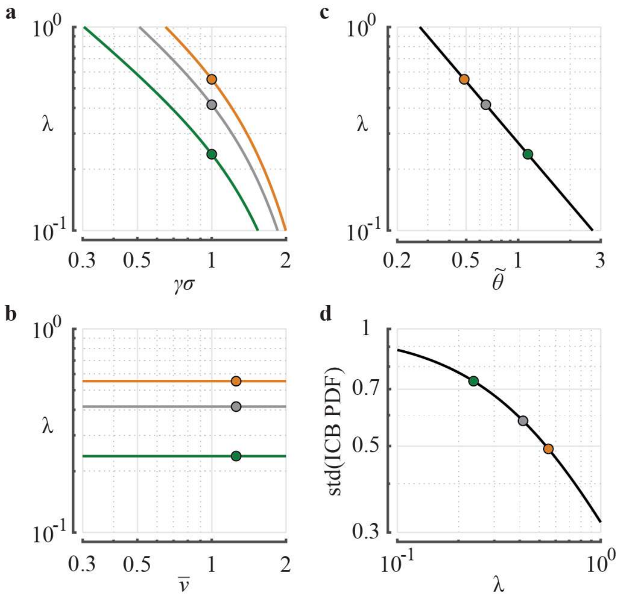


**Figure S4** Poisson model parameters. According to Eq. (5), the shape of the ICBs distribution depends on a single parameter, $\lambda$. Therefore, we can study the effect of changing the model parameters through their effect on $\lambda$. **a**, $\lambda$ vs. $\gamma\sigma$. The three curves denote the three values of $\tilde{\theta}$ depicted in Fig. 4. Orange, $\tilde{\theta} = 0.75 \cdot 0.65$, gray, $\tilde{\theta} = 0.65$ , green, $\tilde{\theta} = 1.75 \cdot 0.65$. Circles denote the value of $\gamma\sigma$ used in Fig. 4, $\gamma\sigma = 1$. $\bar{\nu} = 1.26$ Hz. **b**, $\lambda$ vs. $\tilde{\theta}$, $\gamma\sigma = 1$. $\bar{\nu} = 1.26$ Hz. Circles denote the values of $\tilde{\theta}$ in Fig. 4, color coded as in (a). **c**, $\lambda$ vs. $\bar{\nu}$. The three circles denote the three values of $\tilde{\theta}$ depicted in Fig. 4 as in (a). $\gamma\sigma = 1$. **d**, The width of the ICB distribution vs. $\lambda$. Solid line, the standard deviation of the distribution. Circles denote the values of $\tilde{\theta}$ in Fig. 4, color coded as in (a). $\gamma\sigma = 1$. $\bar{\nu} = 1.26$ Hz. The values of $\lambda$ depicted by circles are $\lambda = 0.24$ (green), $\lambda = 0.41$ (gray) and $\lambda = 0.55$ (orange).

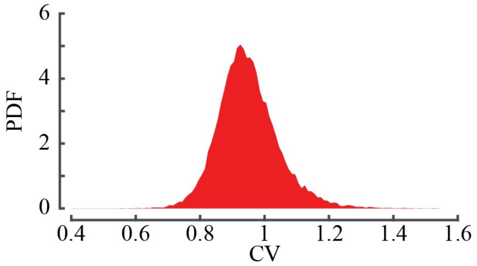


**Figure S5** The distribution of coefficients of variation (CV) of the inter-spike interval distribution of the excitatory neurons in the spiking network model. The CV's were computed over 100 sec of spontaneous activity.

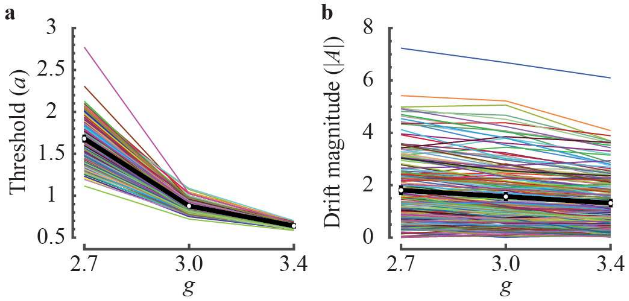


**Figure S6** DDM analysis of the strength of selective inhibition $g$ in the recurrent network model. We fit the 'drift bias' DDM to the responses of the recurrent networks ($n = 200$ networks). For each network in Fig. 6, we computed the posterior-average threshold, $a$, and drift, $A$. **a**, Posterior-average threshold, $a$, for each network (colored thin lines) and their population average (thick black line) are depicted *vs.* $g$. The larger $g$, the smaller is the Posterior-average threshold. **b**, same as in **a** for the absolute value of the drift, $|A|$. Error bars are SEM. Increasing $g$ from 2.7 to 3.4 resulted, on average, in a 62% decrease in the the threshold, $a$, but only in a 27% decrease in the absolute value of the drift. These results indicate that the level of specific inhibition manifests primarily in the decision threshold.

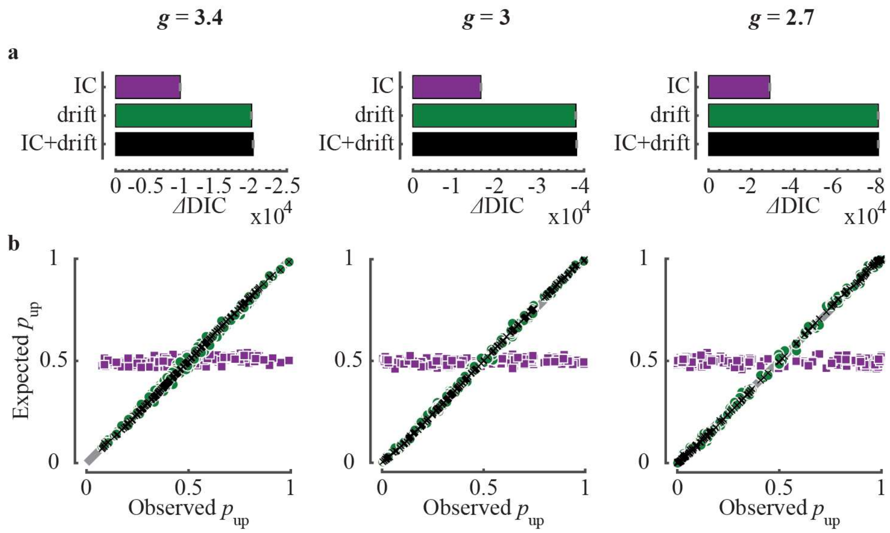


**Figure S7:** DDM analysis of the recurrent spiking network model for the data presented in Fig. 6. **a,** Same analysis as Figs. 1c and 2c. **b**, Same analysis as in Figs. 1d and 2d. Results indicate that similar to the behavioral data, when tested in the framework of the DDM, the ICBs in the recurrent network model are due to drift biases. Best fit orthogonal regression slopes of 'IC bias', 'drift bias' and 'IC+drift bias': $g = 3.4$: 0.05, 0.99, 1.02; $g = 3.0$: 0.02, 1.02, 1.03; $g = 2.7$: 0, 1.01, 1.01.

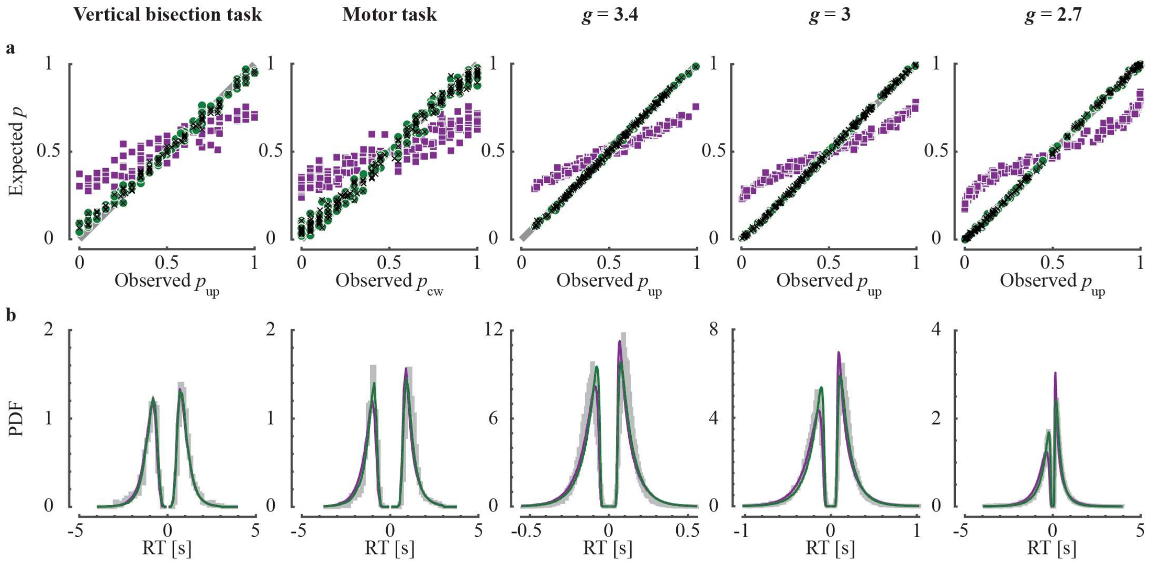


**Figure S8** The 'drift bias' DDM and 'IC+drift bias' DDM but not the 'IC bias' DDM account well for decision-makers' responses. For each dataset, we simulated responses (choices and reaction time) using the corresponding posteriors obtained from the HDDM procedure (2,000 for each participant and 10,000 responses for each recurrent network). **a**, Expected *vs.* observed probability of choice, *p*. Each dot denotes a single decision-maker. Panels from left to right denote the five datasets discussed in the main text: the vertical bisection task, the motor task and the recurrent network model with $g = 3.4$, $g = 3.0$ and $g = 2.7$. Purple squares, 'IC bias' DDM; Green circles, 'drift bias' DDM; Black Xs, 'IC+drift bias' DDM. Gray line, diagonal. Best fit orthogonal regression slopes of 'IC bias', 'drift bias' and 'IC+drift bias': vertical bisection task: 0.45, 0.93, 0.92; motor task: 0.37, 0.95, 0.94; $g = 3.4$: 0.46, 1.03, 1.02; $g = 3.0$: 0.47, 1.03, 1.03; $g = 2.7$: 0.51, 1.01, 1.01. **b**, Reaction-time (RT) distributions. Curves in the positive direction denote the distribution of reaction times for responses congruent with the bias direction. Flipped curves denote the reaction time distribution of those responses that were incongruent with the bias direction. Gray, observed reaction times of the decision-makers; purple, green and black, reaction time distributions of the fitted 'IC bias', 'drift bias' and 'IC+drift bias' DDMs.

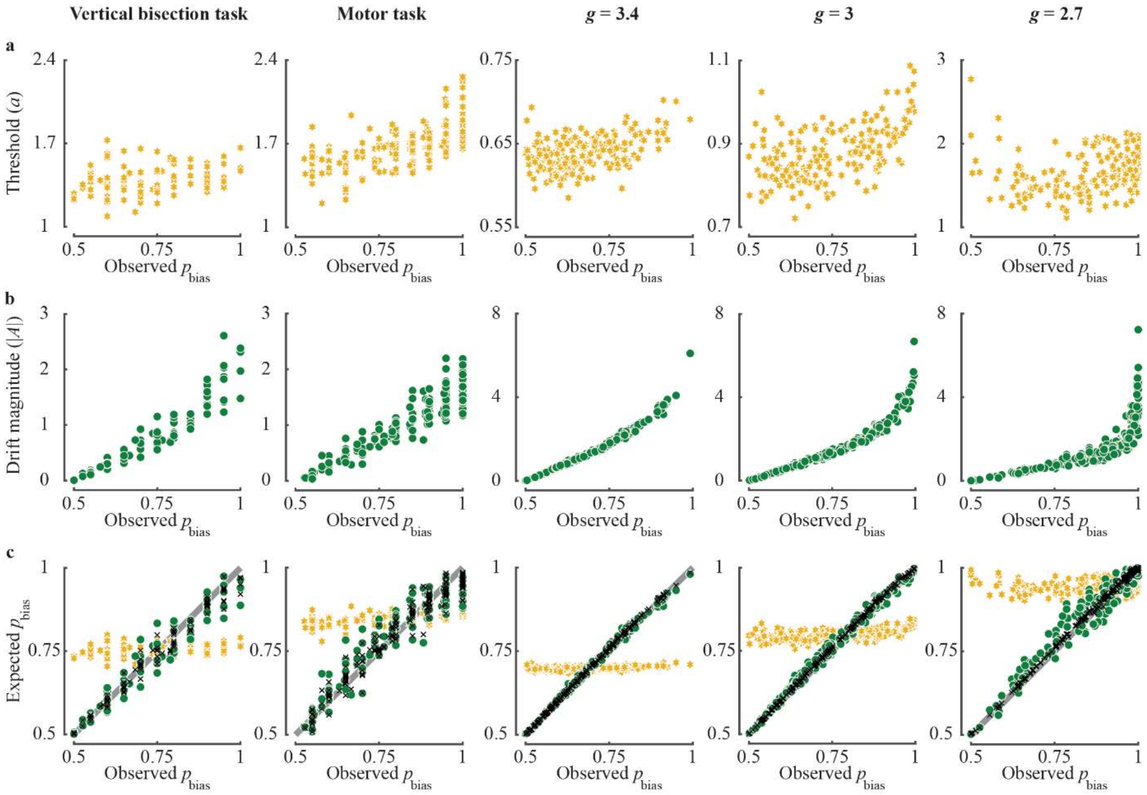


**Figure S9** Contributions of idiosyncratic thresholds and idiosyncratic drift-rates to the ICBs in the ‘drift bias’ DDM. We fitted the ‘drift bias’ DDM to the five datasets. **a**, Posterior-averaged threshold, $a$, and **b**, Posterior-averaged magnitude of the drift, $|A|$, are plotted as a function of the magnitude of the bias, $p_{\text{bias}} = \max\{p, 1-p\}$. Each dot is a single decision-maker and panels from left to right correspond to the five datasets discussed in the main text, the vertical bisection task, the motor task and the recurrent network model with $g = 3.4$, $g = 3.0$ and $g = 2.7$. **c**, We used Eq. (6) with $z = 0.5$ to compute the expected $p_{\text{bias}}$ using the estimated parameters. The DDM-predicted $p_{\text{bias}}$ is plotted for each decision-maker *vs.* the observed $p_{\text{bias}}$. Black Xs: the drift $|A|$ and threshold $a$, are the idiosyncratic estimated ones, as in Fig. S8a. Green circles: the drift is the idiosyncratic estimated one whereas the threshold is its population average. Orange hexagrams: the threshold is idiosyncratic and the drift is its population-average. Gray line is the diagonal. Best fit orthogonal regression slopes of green circles and orange hexagrams: vertical bisection task: 0.93, 0.04; motor task: 0.87, 0.10; $g = 3.4$: 1, 0.02; $g = 3.0$: 0.99, 0.06; $g = 2.7$: 0.96, 0.05. These results indicate that idiosyncrasies in the drift rate rather than in the threshold underlie the ICBs.
